# SMS: Symmetric Mediation Statistics for Powerful High-Dimensional Mediation Analysis

**DOI:** 10.64898/2026.06.10.730748

**Authors:** Yi Wang, Shiyu Yan, Hai-Jun Wang, Yi-Juan Hu

## Abstract

**Background:** Mediation analysis of high-dimensional features, particularly molecular-level omics features, provides important opportunities to uncover biological mechanisms underlying human health and disease. However, two central statistical challenges remain: testing the composite-null hypothesis and maintaining power when the exposure–mediator and mediator–outcome associations differ substantially in statistical significance. Existing methods typically rely on accurate estimation of the proportions of the three null types or on the maximum of the two association *p*-values, and may not always control the FDR well and may have limited power under imbalanced significance.

**Methods:** We propose SMS, a new statistical framework based on symmetric mediation statistics. By exploiting symmetry, SMS calibrates the composite null distribution as a whole for FDR control. It also allows flexible combinations of the two association *p*-values, including the maximum, and then enables construction of an omnibus test. Moreover, it permits direct use of effect-size estimates, bypassing the need to compute *p*-values.

**Results:** SMS controlled the FDR across a wide range of simulation scenarios while achieving a substantial sensitivity gain, often around 20 percentage points, over existing methods including HDMT, DACT, and DEI-B. Applications to a metabolomics dataset and a DNA methylation dataset further corroborated these findings. Notably, SMS discovered five plausible mediators in the metabolomics dataset that were missed by all existing methods considered.

## Introduction

In recent years, technologies for profiling diverse omics layers, including transcriptomics, epigenomics, metabolomics, and metagenomics, have advanced rapidly. The resulting high-dimensional molecular data provide new opportunities to investigate biological mechanisms underlying human health and disease. One mechanism of particular interest is causal mediation, in which an exposure affects an outcome through intermediate variables.

As a motivating example, we considered the Peking University–Shandong Preschool Childhood Obesity case-control study (PKU-SPCO Study) [1]. Conducted by the Peking University research team, the study recruited 128 preschool children from Shandong Province, China, to investigate lifestyle, gut microbiome, and fecal metabolomic factors associated with childhood obesity. Prior studies [1–4] have established a strong association between maternal obesity and childhood obesity, which was also observed in our data (odds ratio = 5.5; chi-squared *p*-value = 2.1 × 10^−5^). Our goal was to identify metabolites that mediate the effect of maternal obesity on childhood obesity. To this end, we performed untargeted metabolomics profiling using mass spectrometry, yielding measurements of 1,152 metabolites.

Beyond high dimensionality, a central challenge in causal mediation analysis is the composite nature of the null hypothesis. Let *E* denote the exposure, *M* = (*M*_1_, …, *M*_*J*_) the *J* features, *O* the outcome, and *C* the confounding covariates. The mediation relationships under this notation are illustrated in Figure 1(a). For feature *j* to have a mediation effect, both the exposure–mediator (E–M) and mediator–outcome (M–O, conditional on the exposure) associations must be present. Therefore, the null hypothesis of no mediation for feature *j* is composite and comprises three types:

**Figure 1:**
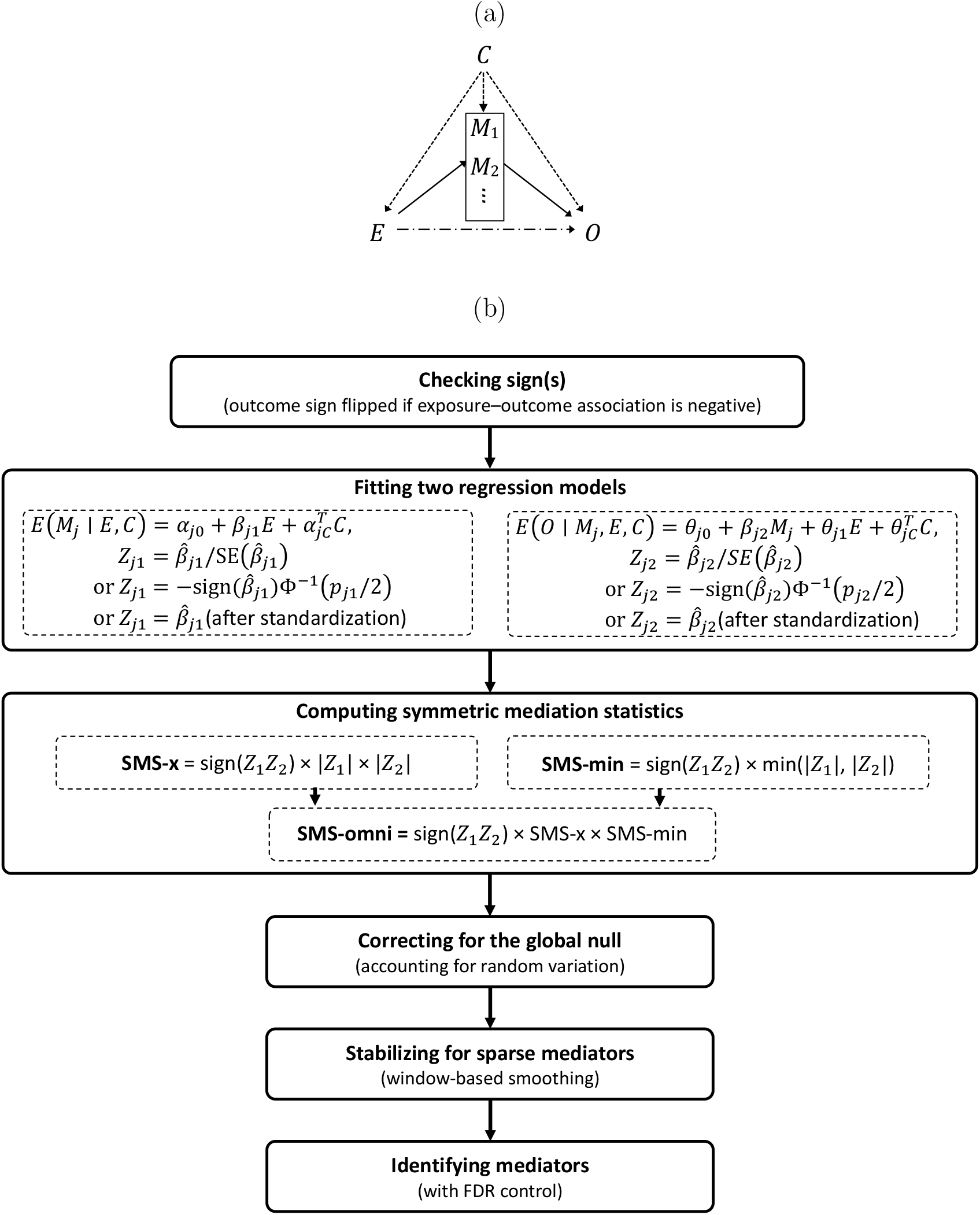
(a) Directed acyclic graph (DAG) for mediation analysis. *E* denotes the exposure, *M*_1_, *M*_2_, … denote candidate mediators, *O* denotes the outcome, and *C* denotes measured confounders. Solid arrows indicate mediation pathways, the dash-dotted arrow indicates the direct effect, and dashed arrows indicate confounding effects. (b) Workflow of the SMS framework.

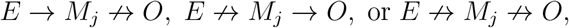

referred to as type-I, type-II, and type-III nulls, respectively.

While earlier methods such as Sobel’s test [5] and the MaxP test [6] ignored the composite-null structure of mediation analysis, more recent approaches explicitly account for it. HDMT [7] uses the maximum of the E–M and M–O association *p*-values as the test statistic and models its null distribution as a mixture of distributions corresponding to the three null types. The mixing proportions, which equal the proportions of the three null types, are estimated using an adaptation of the procedure of [8]. DACT [9] instead constructs a test statistic from a mixture of null-type-specific *p*-values, one of which is based on the maximum *p*-value, with the null proportions estimated following Jin and Cai (2007) [10]. When one null type dominates, DACT adopts a *U* (0, 1) null distribution; otherwise, it estimates the null distribution using Efron’s empirical null framework [11]. Thus, both HDMT and DACT depend critically on accurate estimation of the null proportions, which can be difficult in practice [12]. Most recently, DEI-B [13] also uses the maximum *p*-value as the test statistic, but avoids estimating the null proportions by deriving a closed-form *p*-value for the composite null as a whole. However, this closed-form result is derived under the assumption of a very small proportion of true mediators and therefore tends to be conservative otherwise.

We applied HDMT, DACT, and DEI-B to the PKU-SPCO metabolomics data to identify metabolic mediators, but none detected any signals, despite strong biological evidence reported in the literature [14–17]. This lack of detection may be explained by two factors. First, the null distributions of the test statistics may not be well calibrated, leading to overly conservative results. Second, and likely more importantly, these methods rely on the maximum *p*-value, which effectively requires both the E–M and M–O associations to be significant for a mediator to be detected. As a result, they can lose power when the two associations differ substantially in significance level, a situation commonly encountered in practice. For example, in our PKU-SPCO metabolomics study, maternal obesity exhibits weaker associations with childhood metabolism than childhood obesity does (Figure 2, left panel).

**Figure 2:**
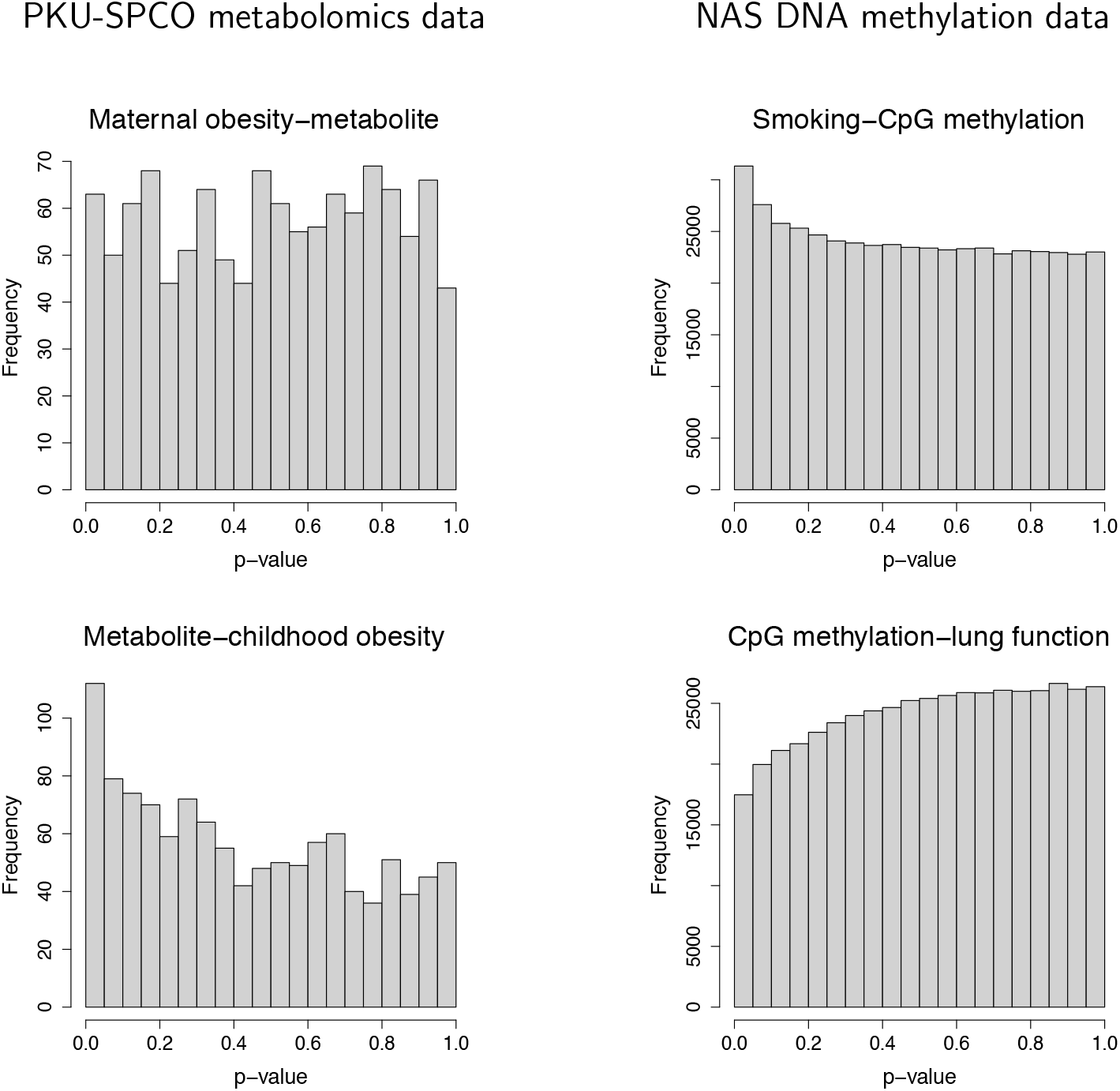
Distributions of *p*-values for the E–M associations (upper panel) and M–O associations (lower panel) in the two real datasets.

Moreover, HDMT, DACT, and DEI-B all require pairs of *p*-values from the E–M and M–O association tests. In high-dimensional settings, however, obtaining valid *p*-values can be difficult. For example, methods such as LDM [18] and LOCOM [19] rely on resampling procedures, which become computationally intensive when estimating extremely small *p*-values. In recent years, methods that bypass *p*-values to achieve false discovery rate (FDR) control, including the knockoff [20, 21] and data-splitting procedures [22], have received considerable attention. Yet these ideas have not yet been incorporated into high-dimensional mediation analysis.

In this article, we introduce SMS, a new framework for mediation analysis with high-dimensional mediators based on symmetric mediation statistics. The proposed statistics are designed to (1) calibrate the rejection threshold for FDR control under the composite null as a whole without estimating the proportions of its component null types, (2) accommodate imbalance in the significance of the E–M and M–O associations, and (3) allow direct use of effect-size estimates while bypassing *p*-value calculation. The complete workflow of SMS is illustrated in Figure 1(b), and its key features relative to existing methods are summarized in Table 1. We evaluate the performance of SMS through extensive simulation studies and further apply it to two real datasets. In addition to the PKU-SPCO metabolomics data, we reanalyzed DNA methylation data from the Normative Aging Study (NAS) [23], which was previously examined by DACT and DEI-B, thereby enabling direct comparison with these methods.

**Table 1:** High-dimensional mediation analysis methods evaluated in this study.

| Method | Reference | Require<br>null<br>proportions | Use<br>maximum<br>$p$-value | Require<br>association<br>$p$-values | Rely on<br>the BH<br>procedure |
| --- | --- | --- | --- | --- | --- |
| HDMT | Dai et al., 2022 | * | * | * |  |
| DACT | Liu et al., 2022 | * | * | * | * |
| DEI-B | Liu, 2025 |  | * | * | * |
| SMS | This work |  |  |  |  |

Note: \* denotes a limiting feature of the method. BH stands for Benjamini–Hochberg [28].

## Methods

### Symmetric mediation statistic (SMS)

We begin with a simple setting in which the exposure and outcome are univariate, and both the high-dimensional features and the outcome are continuous; this setting is generalized later. We assume that all features have been standardized to have equal variances, although we show later that one of the proposed statistics is invariant to feature scaling. Following the modern causal inference framework [24], we further assume no unmeasured confounders and no exposure–mediator or mediator–mediator interactions. The classical multiple-mediator model [25] specifies a linear model for each mediator and a linear model for the outcome that includes the effects of all mediators:

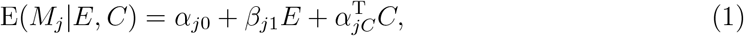

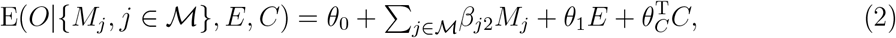

where ℳ denotes the index set of true mediators, *β*_*j*1_ characterizes the effect of the exposure *E* on the *j*th feature *M*_*j*_ conditional on the confounding covariates *C*, and *β*_*j*2_ characterizes the effect of *M*_*j*_ on the outcome *O* conditional on *E* and *C* and the remaining mediators. Under this model, the overall (or total) mediation effect is given by _*j* ∈ℳ_ *β*_*j*1_*β*_*j*2_ [25]. If the mediators are mutually independent conditional on *E* and *C*, each product *β*_*j*1_*β*_*j*2_ can be interpreted as the mediation effect through the individual mediator *M*_*j*_. Even without this conditional independence assumption, a non-zero value of *β*_*j*1_*β*_*j*2_ indicates a marginal mediation signal. Therefore, mediator detection can be formulated as testing whether *β*_*j*1_*β*_*j*2_ = 0 for each feature *M*_*j*_.

Following [7, 9, 13], we estimate *β*_*j*1_ from (1) but estimate *β*_*j*2_ by replacing the joint outcome model in (2) with a marginal outcome model for each *M*_*j*_:

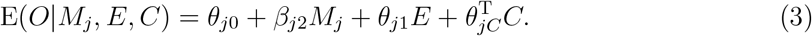

The corresponding estimates are denoted by 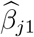 and 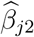. We then construct the following product-form statistic for testing the mediation effect of *M*_*j*_:

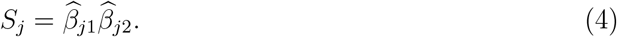

This is the same product statistic used in Sobel’s test, where inference proceeds by estimating the standard error of *S*_*j*_ and constructing a Wald-type test statistic. However, this test is asymptotically valid only under the type-I and type-II nulls, conservative in finite samples under these nulls, and especially conservative under the type-III null [9]. Instead, we exploit an important symmetry property of *S*_*j*_ under the composite-null hypothesis of no mediation. This symmetry forms the basis for mediator detection with FDR control, in a spirit similar to the knockoff and data-splitting procedures.

The symmetry property follows from two key observations. First, under the null hypothesis of no E–M association or no M–O association, 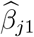 or 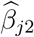 is symmetric about zero, respectively. Second, the two coefficient estimates, 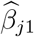 and 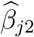, are independent, as shown by [9]. Therefore, under the composite null, where at least one of the two coefficient estimates is symmetric about zero, their product *S*_*j*_ is also symmetric about zero.

In contrast to this null symmetry, under the mediation alternative, the collection of *S*_*j*_ statistics corresponding to true mediators tends to align in a common direction. Without loss of generality, we orient the outcome so that the E–O association is positive, reversing its sign when necessary. Under a positive total effect of the exposure on the outcome, most mediator-specific effects are expected to be positive. Oppositely directed effects may occur but are likely uncommon; therefore, we focus on mediators whose effects are aligned with the total effect. This implies that, for a true mediator, the E–M and M–O associations are expected to have the same sign, either both positive or both negative. This sign pattern is supported by the NAS data analysis results (Table S1), where all mediators identified by existing methods, which do not impose this assumption, satisfy this pattern.

Under this common-direction assumption, a larger positive value of *S*_*j*_ provides stronger evidence that feature *M*_*j*_ is a mediator. This ranking is meaningful because *M*_*j*_’s are standardized to have equal variances, making the *β*_*j*1_’s directly comparable across features, and likewise for the *β*_*j*2_’s. Note that *β*_*j*1_ and *β*_*j*2_ for a given feature need not be on the same scale.

The distributional properties of *S*_*j*_ discussed above can be empirically verified in the two real datasets. As shown in Figure 3 (upper panel), the distribution of *S*_*j*_ in both datasets exhibits two clear components: a large component that is approximately symmetric about zero, corresponding to null features, and a small component in the positive tail, corresponding to plausible mediators. Because this statistic is analogous to the “symmetric statistic” or “mirror statistic” proposed by Dai et al. [22], we refer to *S*_*j*_ as the symmetric mediation statistic (SMS).

**Figure 3:**
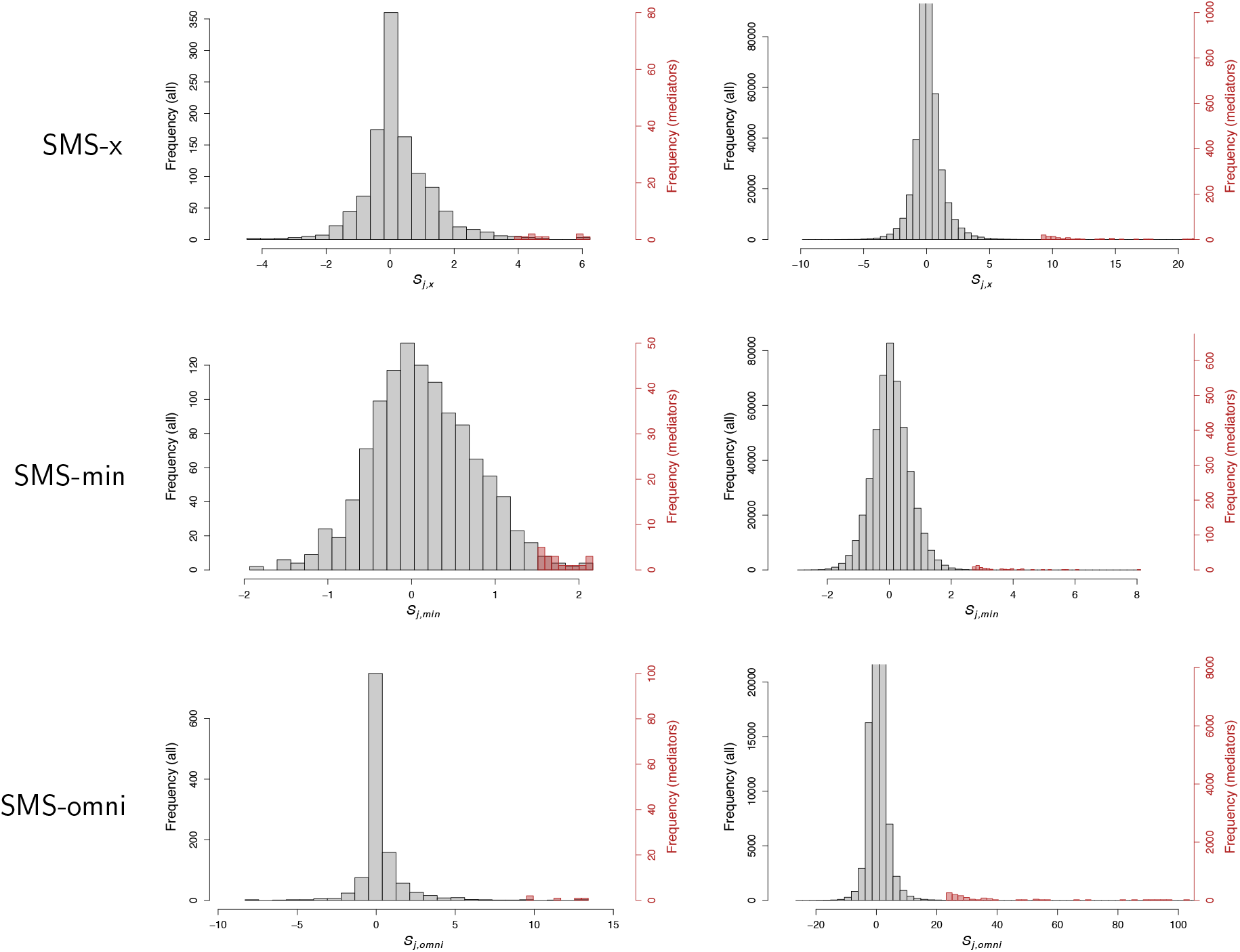
Distributions of the statistics in SMS-x, SMS-min, and SMS-omni, for all features in the two real datasets. The red region denotes mediators identified by the corresponding SMS method and is enlarged to show greater detail. For the NAS dataset, the x-axis is truncated at 20 for SMS-x and at 100 for SMS-omni, and the y-axis is truncated at 8,000 and 2,000, respectively, to show greater detail in the positive tail of the gray region.

Next, we use the symmetry of the SMS among null features to obtain an upper bound on the number of false positives. Specifically, let 𝒩 denote the index set of null features. For any *t >* 0,

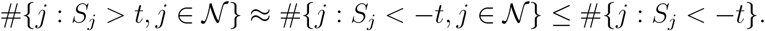

Thus, the false discovery proportion (FDP),

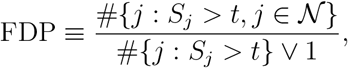

is upper bounded by

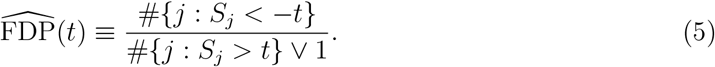

For a nominal FDR level *α*, we define the data-driven threshold,

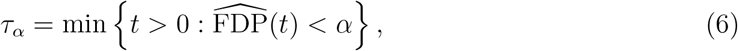

and identify mediators as those features satisfying *S*_*j*_ *> τ*_*α*_.

To determine the threshold, 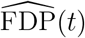 is evaluated over {*t*_1_, *t*_2_, …, *t*_*J*_}, the increasingly ordered values of {|*S*_1_|, |*S*_2_|, …, |*S*_*J*_ |}. Because 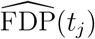 generally decreases as *t*_*j*_ increases in the threshold region, *τ*_*α*_ is chosen as the first *t*_*j*_ at which 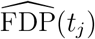 falls strictly below the nominal FDR level (Figure S1). As shown in [22], this procedure controls the FDR asymptotically. By comparing the symmetric distribution of null features as a whole with the positive tail enriched for mediators, our method avoids estimating the proportions of different null types separately. In this respect, it shares a key advantage with DEI-B.

### 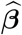-value vs. *Z*-score

When standard error estimates SE 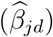, *d* = 1, 2, are available, the effect-size estimates 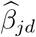 used in *S*_*j*_ can be replaced by the corresponding *Z*-scores, 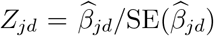. Equivalently, if the corresponding two-sided *p*-values *p*_*jd*_ are available, the *Z*-scores can be recovered as 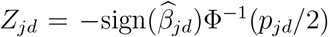, where ^Φ^(*·*) is the cumulative distribution function of the standard normal distribution. This standardization eliminates the need to standardize features beforehand. We therefore generally recommend using the *Z*-score-based statistic

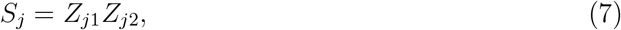

which is invariant to variable scaling and more directly reflects the statistical significance of the two associations. Nevertheless, as shown below, the two statistics, (4) and (7), are closely related and typically exhibit very similar empirical performance.

The two standard error estimates take the forms:

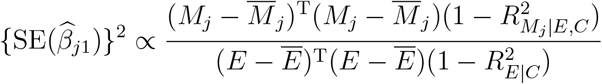

and

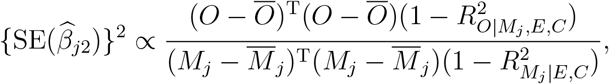

where 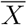 denotes the mean of a generic vector *X*, 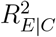 is the coefficient of determination 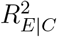 from the regression of *E* on *C*, and 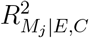 and 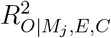 are coefficients of determination from models (1) and (3), respectively. Note that coefficients of determination are invariant to variable scaling. After ignoring feature-independent factors, the *Z*-score-based SMS in (7) differs from the 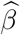-based SMS in (4) only by a factor of 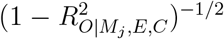. In practice, 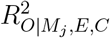 is usually small for each feature *M*_*j*_, because any individual feature typically explains only a small fraction of the variation in *O*. Thus, the *Z*-score-based SMS is expected to perform similarly to the 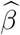-based SMS.

An interesting result, perhaps counterintuitive at first, is that the 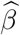-based SMS in (4) is invariant to feature scaling and therefore does not require prior feature standardization. This invariance arises because rescaling *M*_*j*_ changes 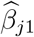 and 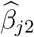 in opposite directions, so the feature scale factors cancel in the product 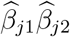. Equivalently, the same cancellation appears in the product of the standard errors,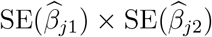. However, this property holds only for the product SMS, and does not necessarily extend to other forms of SMS discussed below. To guard against such cases, we standardize features by default whenever the 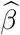-based SMS is used.

### Minimum SMS and omnibus SMS

For the product SMS in (7), the distributions of type-I, type-II, and type-III null statistics are all symmetric about zero, but they may have different variances. In particular, type-I and type-II null statistics tend to have larger variances than type-III null statistics (Figure S2, left panel); asymptotically, their variance is four times that of type-III null statistics [9]. When type-I and/or type-II nulls make up a non-negligible fraction of all null features, they can dominate the negative-tail calibration and thereby largely determine the threshold *τ*_*α*_, whereas type-III nulls, which typically form the largest cluster, contribute little to this calibration. When true mediators are also sparse, the resulting small alternative cluster can be difficult to distinguish from the small type-I/type-II null cluster, resulting in reduced power.

To address this issue, we consider the minimum SMS as the signed minimum of |*Z*_*j*1_| and |*Z*_*j*2_|:

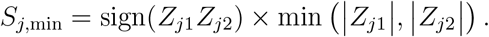

Under this form, the distributions of the three types of null statistics have similar variances (Figure S2, right panel), jointly forming a clearer contrast with the alternative cluster and thereby improving power. For this SMS, using 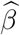 values is less appropriate than for the product SMS (4), even with feature standardization, because the two regression coefficients represent different effects and may have different scales. As a result, min 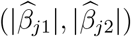 may primarily reflect the coefficient on the smaller scale rather than the weaker association. We therefore define this SMS using *Z*-scores whenever possible. It can then be used to identify mediators through the same procedure as in (5) and (6). To distinguish this statistic from the product statistic, we rewrite the latter in (7) as

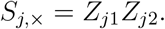

We refer to the tests based on *S*_*j*,×_ and *S*_*j*,min_ as SMS-x and SMS-min, respectively.

In practice, it is not known a priori which statistic will be more powerful. We therefore construct an omnibus test that combines the information from *S*_*j*,×_ and *S*_*j*,min_, aiming to achieve performance close to the better of the two tests. Specifically, we define the omnibus SMS as

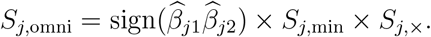

This statistic is again symmetric about zero, with larger positive values indicating stronger evidence for mediation. Although *S*_*j*,min_ and *S*_*j*,×_ are not independent, the sign 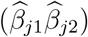 term preserves the sign structure and hence the symmetry of *S*_*j*,omni_. The same FDP estimation and thresholding procedure in (5) and (6) can then be applied to identify mediators. We refer to this omnibus test as SMS-omni.

In the data-splitting framework [22], the symmetric statistic defined as the sum of two coefficient estimates was shown analytically to be optimal relative to the product and minimum statistics. This result, however, does not appear to carry over to our setting (results not shown), primarily because our inferential target is different. The data-splitting framework aims to identify features for which two estimates of the same effect size, obtained from split datasets, are both large. In contrast, mediation analysis targets features with large mediation effects, *β*_*j*1_*β*_*j*2_, allowing one association to be very large even when the other is only moderately large.

### Correction under the global-null setting

The negative-tail calibration procedure in (5) and (6) performs well when sufficiently many true signals are present in the positive tail, as required by [22]. Under the global null, when signals are completely absent, random variation yields − min_*j*_ *S*_*j*_ *<* max_*j*_ *S*_*j*_ with 50% probability, where *S*_*j*_ denotes any of *S*_*j*,×_, *S*_*j*,min_, or *S*_*j*,omni_. In this case, evaluating 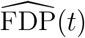 at *t* = − min_*j*_ *S*_*j*_ gives 0, so the inequality in (6) is satisfied and *τ*_*α*_ *≤* − min_*j*_ *S*_*j*_. Consequently, the detection set {*j* : *S*_*j*_ *> τ*_*α*_} is nonempty and contains at least the feature corresponding to max_*j*_ *S*_*j*_ (Figure S1, left). Under the global null, this yields an empirical FDP of 1 and, after accounting for its 50% probability, an empirical FDR of approximately 50% (Figure S3).

Even when true signals are present, if they are sparse, the same phenomenon can still lead to large empirical FDR values.

This problem occurs when random variation drives the negative-tail count (i.e., numerator) of 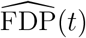 to zero, making the criterion in (6) too easy to satisfy and leading to overly aggressive detection. To reduce the effect of this randomness, we add a correction term, *αC*(*t*), to the negative-tail count of 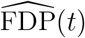 whenever it becomes zero, where *α* is the nominal FDR level and *C*(*t*) is a binomial quantile specified below. Specifically, we define the corrected FDP estimator as

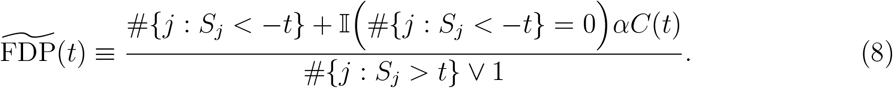

To motivate the choice of *C*(*t*), suppose that 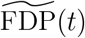 is monotonically decreasing in *t* in the threshold region. Under the global null, symmetry implies that the two events − min_*j*_ *S*_*j*_ *≥* max_*j*_ *S*_*j*_ and − min_*j*_ *S*_*j*_ *<* max_*j*_ *S*_*j*_ each occur with probability approximately 50%. When − min_*j*_ *S*_*j*_ *≥* max_*j*_ *S*_*j*_, for any *t ≥* max_*j*_ *S*_*j*_, we have #{*j* : *S*_*j*_ *> t*} = 0 and 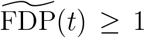; hence, no detection is made and the empirical FDP is 0. When − min_*j*_ *S*_*j*_ *<* max_*j*_ *S*_*j*_, taking *t* = − min_*j*_ *S*_*j*_ gives #{*j* : *S*_*j*_ *<* −*t*} = 0 and 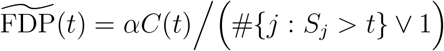. If *C*(*t*) is chosen such that

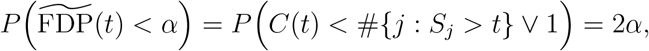

then, because this case occurs with probability approximately 50%, the empirical FDR is approximately controlled at level *α*.

We next describe how to determine *C*(*t*) at *t* = − min_*j*_ *S*_*j*_. Under the global null, #{*j* : *S*_*j*_ *> t*} follows a binomial distribution with *J* trials and success probability

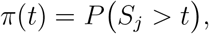

where *P* (.) is evaluated under the composite-null distribution. If *π*(*t*) were known, *C*(*t*) could be defined as the (1 − 2*α*)-quantile of this binomial distribution, or equivalently, of the distribution of #{*j* : *S*_*j*_ *> t*} V 1, after left truncation at 1. A natural approach is to estimate *π*(*t*) from the left tail of the empirical distribution of *S*_*j*_. However, because *π*(*t*) is typically extremely small at *t* = − min_*j*_ *S*_*j*_, this empirical estimate may be imprecise when the number of features is moderate, for example *J* = 1,000.

To obtain a smoother empirical distribution of *S*_*j*_, we generate augmented statistics 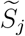 by independently resampling one value, 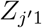, from {*Z*_*j*1_, *j* = 1, …, *J*} and one value, 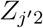, from {*Z*_*j*2_, *j* = 1, …, *J*}, and then combining them in the same way as the original statistic. We repeat this procedure to generate 100,000 augmented 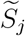 values. We then estimate *π*(*t*) as the empirical proportion of 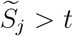, or preferably, as one half of the empirical proportion of 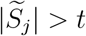. This augmented empirical distribution provides a stable approximation to the composite-null distribution, while avoiding explicit estimation of the proportions of different null types. When the type-I and type-II nulls do not coexist, although either may coexist with type-III nulls, the augmented empirical distribution exactly recovers the composite-null distribution. When type-I and type-II nulls coexist, however, a *Z*_*j*1_ value reflecting an E–M association may be paired by chance with a 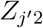 value reflecting an M–O association, producing an alternative-like 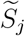 and leading to overestimation of *π*(*t*). In practice, this overestimation should be small: if type-I and type-II nulls each account for 10% of the features, only 1% of augmented statistics arise from such mismatched pairs. Moreover, overestimation of *π*(*t*) leads to overestimation of *C*(*t*) when calibrating the tail probability at 2*α*, making the empirical FDR conservative rather than anti-conservative.

When mediation signals are present but sparse, for example with only 10 true mediators, 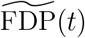 may not fall below *α* until *t* = − min_*j*_ *S*_*j*_ (Figure S1, middle), where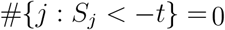. Thus, the correction can still influence the thresholding rule. In contrast, when mediators are dense, #{*j* : *S*_*j*_ *<* −*t*} *>* 0 typically holds at the selected threshold *t* = *τ*_*α*_ (Figure S1, right), so the correction has little or no effect.

A related conservative alternative is the “plus-one” FDP estimator of Barber and Candès [20], which adds one to the negative-tail count of 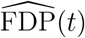:

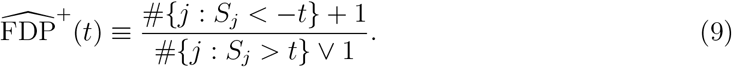

This “plus-one” correction is considerably more conservative than our adaptive correction in 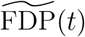 and can substantially reduce power in sparse-mediator settings, a pattern confirmed in our simulation studies (Figure S4).

### Stabilization under the sparse-mediator setting

In sparse-mediator settings, 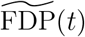 can fluctuate rather than decrease smoothly with increasing *t*, which may yield an unstable threshold *τ*_*α*_ and overly aggressive detection (Figure S1, middle). To stabilize threshold selection, we smooth 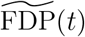 over *t* using the local moving-average procedure described below.

Let {*t*_1_, *t*_2_, …, *t*_*J*_} be the ordered values of {|*S*_1_|, |*S*_2_|, …, |*S*_*J*_ |}, and let *W* denote the one-sided window size. For each *t*_*j*_, we smooth 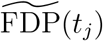 by averaging it over a local window containing up to *W* neighbors on either side:

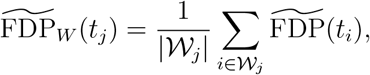

where *𝒲*_*j*_ = {*i* : max(1, *j* − *W*) *≤ i ≤* min(*J, j* + *W*)} is the index set of local neighbors and |*𝒲*_*j*_| denotes its cardinality. The threshold *τ*_*α*_ is then chosen as the smallest *t*_*j*_ such that 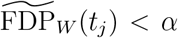. Our numerical studies suggest that a small smoothing window, *W* = 2, is sufficient to control the FDR (Figure S3). We therefore set *W* = 2 as the default window size for all SMS implementations. Similar to the correction procedure described above, the stabilization procedure has little effect when mediators are dense (Figure S1, right).

More broadly, our work appears to be the first to systematically examine and address FDR-control challenges in the global-null and sparse-signal settings for procedures based on symmetric statistics, including knockoff- and data-splitting-based methods. The proposed correction and stabilization procedures therefore extend beyond our SMS mediation analysis and may be useful for a broader class of FDR-control methods built on symmetric statistics.

### Extensions to other settings

The SMS framework can also accommodate non-continuous outcomes, such as binary or time-to-event outcomes. In these settings, equation (3) can be replaced by an appropriate model, such as a generalized linear model (GLM) or a survival model. When the outcome model is nonlinear, fitting a feature-specific marginal model as in (3) may attenuate effect-size estimates [26]. This attenuation, however, may affect power but not error-rate control.

In some applications, each feature may be paired with its own outcome. For example, Dai et al. [7] considered mediation triplets by identifying CpG methylation sites within 500 kb of each risk single-nucleotide polymorphism (SNP) and linking the DNA methylation levels at these sites to the expression levels of the corresponding genes, resulting in 69,902 triplets for mediation analysis. In this setting, each outcome should be oriented, by reversing its sign when necessary, so that the corresponding E–O association is positive.

Whenever available, *Z*-scores or *p*-values are preferred. When only effect-size estimates are available, however, they can be used directly to construct the product SMS without prior feature standardization. This is because the feature-scale invariance of the 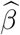-based product SMS extends to all settings described above. To see this, consider first a general univariate outcome and a feature-specific outcome model, such as a GLM or Cox model, that is related to feature *M*_*j*_ through the term *β*_*j*2_*M*_*j*_. The coefficient estimate 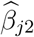 is thus inversely related to the scale of *M*_*j*_. Meanwhile, the estimate 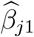 from model (1) is proportional to the scale of *M*_*j*_. The two scale factors of *M*_*j*_ cancel in the product of 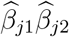. When the outcome is feature-specific, 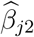also depends on the scale of outcome *j*. After 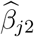 is rescaled by the empirical standard deviation of outcome *j*, prior feature standardization remains unnecessary for the 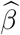-based product SMS. By contrast, feature standardization is always required for the 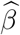-based minimum SMS.

## Results

### Simulation studies

We conducted extensive simulation studies to evaluate the proposed methods, SMS-omni, SMS-x, and SMS-min, relative to existing methods, including HDMT, DACT, and DEI-B. Motivated by the two real datasets, PKU-SPCO and NAS, we varied the sample size *n* from 100 to 500 and the feature dimension *J* from 1,000 to 50,000.

For each feature dimension *J*, we considered a global-null setting with no mediator, sparse-mediator settings with 10 or 25 mediators, and dense-mediator settings with 50 or 100 mediators. We further considered four composite-null configurations, with the numbers of type-I and type-II null features set to (10, 10), (100, 10), (10, 100), and (100, 100), denoted by SS, DS, SD, and DD, respectively, where S and D indicate sparse and dense type-I or type-II nulls. All remaining null features were type-III nulls.

For each simulation replicate, we generated a binary exposure *E*_*i*_ ∈ {0, 1}, with equal numbers of subjects assigned to the two exposure groups. We then generated *J* features according to *M*_*ij*_ = *β*_*j*1_*E*_*i*_ + *ε*_*ij*_, where *ε*_*ij*_ ~ *N* (0, 1). Continuous outcomes were generated as 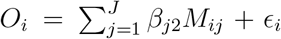, where *ϵ*_*i*_ ~ *N* (0, 1). Binary outcomes were sampled as *O*_*i*_ ~ Bernoulli(*π*_*i*_), with log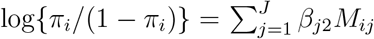. In settings where each mediator had its own outcome, denoted by *O*_*ij*_, the outcome was generated as *O*_*ij*_ = *β*_*j*2_*M*_*ij*_ + *ϵ*_*i*_.

The coefficients *β*_*j*1_ and *β*_*j*2_ were set to zero according to the corresponding null type; otherwise, they were sampled as *β*_*j*1_ ~ *c*_1_Beta(2, 5) and *β*_*j*2_ ~ *c*_2_Beta(2, 5). The scaling parameters (*c*_1_, *c*_2_) were initially set to (1, 1) and then varied over (0.25, 4), (0.5, 2), (2, 0.5), and (4, 0.25) to induce different relative strengths of the E–M and M–O associations. Note that the Beta(2, 5) distribution is asymmetric on [0, 1], has mode 0.2, and yields only positive values. To allow both positive and negative effects, one-third of the nonzero coefficients were randomly assigned negative signs. For true mediators, the signs of *β*_*j*1_ and *β*_*j*2_ were assigned jointly to ensure positive mediation effects.

For each simulated dataset, we obtained effect-size estimates and *p*-values by fitting a linear regression of *M*_*ij*_ on *E*_*i*_, followed by either a linear or logistic regression of *O*_*i*_ or *O*_*ij*_ on *M*_*ij*_ and *E*_*i*_, depending on the outcome type. HDMT, DACT, and DEI-B were applied directly to the resulting *p*-values. To ensure comparability with these methods, we first implemented the SMS methods using *Z*-scores, and then evaluated an alternative implementation based on effectsize estimates after feature standardization. For each method, empirical FDR and sensitivity were assessed at the nominal FDR level of 0.2. We used 1,000 simulation replicates for the global-null and sparse-mediator settings to ensure stable FDR estimates, and 100 replicates for the dense-mediator settings.

### Simulation results

We mainly present results for a continuous outcome, unless explicitly stated otherwise, because the results for a binary outcome and feature-specific outcomes showed similar patterns. We first examined whether the proposed correction and stabilization procedures were necessary for the SMS methods to control the FDR. Figure S3 shows that, without either correction or stabilization, the empirical FDR reached approximately 0.5 under the global null and remained substantially inflated when 10 mediators were present. The correction procedure alone reduced the FDR to the nominal level under the global null, but some inflation persisted in the 10-mediator setting. Applying stabilization with *W* = 1 reduced this remaining inflation, whereas using *W* = 2 eliminated it entirely. In settings with dense mediators, the combined correction and stabilization procedures had little impact on the results (results not shown). We also found that the “plus-one” correction in (9) was overly conservative. Figure S4 showed that this correction substantially reduced sensitivity, particularly in sparse-mediator settings, with a loss of up to 20 percentage points. Therefore, all subsequent SMS results were obtained using both the proposed correction and stabilization procedures.

Figures 4, 5, and 6 summarize the empirical FDR and sensitivity results for *J* = 1,000, 5,000, and 50,000, respectively. The SMS methods, including SMS-omni, SMS-x, and SMS-min, consistently maintained the FDR below the nominal level across all settings. The FDR was conservative under the global null across all composite-null configurations because of the stabilization strategy. It was also conservative in the DD setting with sparse mediators due to overestimation of *π*(*t*). DEI-B yielded empirical FDR close to the nominal level in the global-null and sparse-mediator settings, but became conservative as the number of true mediators increased, particularly when *J* was relatively small (*J* = 1,000). HDMT was generally conservative when *J* was not large, although it occasionally showed slight inflation under the global null, for example when *J* = 1,000 under SS. Interestingly, the FDR of DACT was substantially inflated when *J* = 1,000 but markedly deflated when *J* = 50,000.

**Figure 4:**
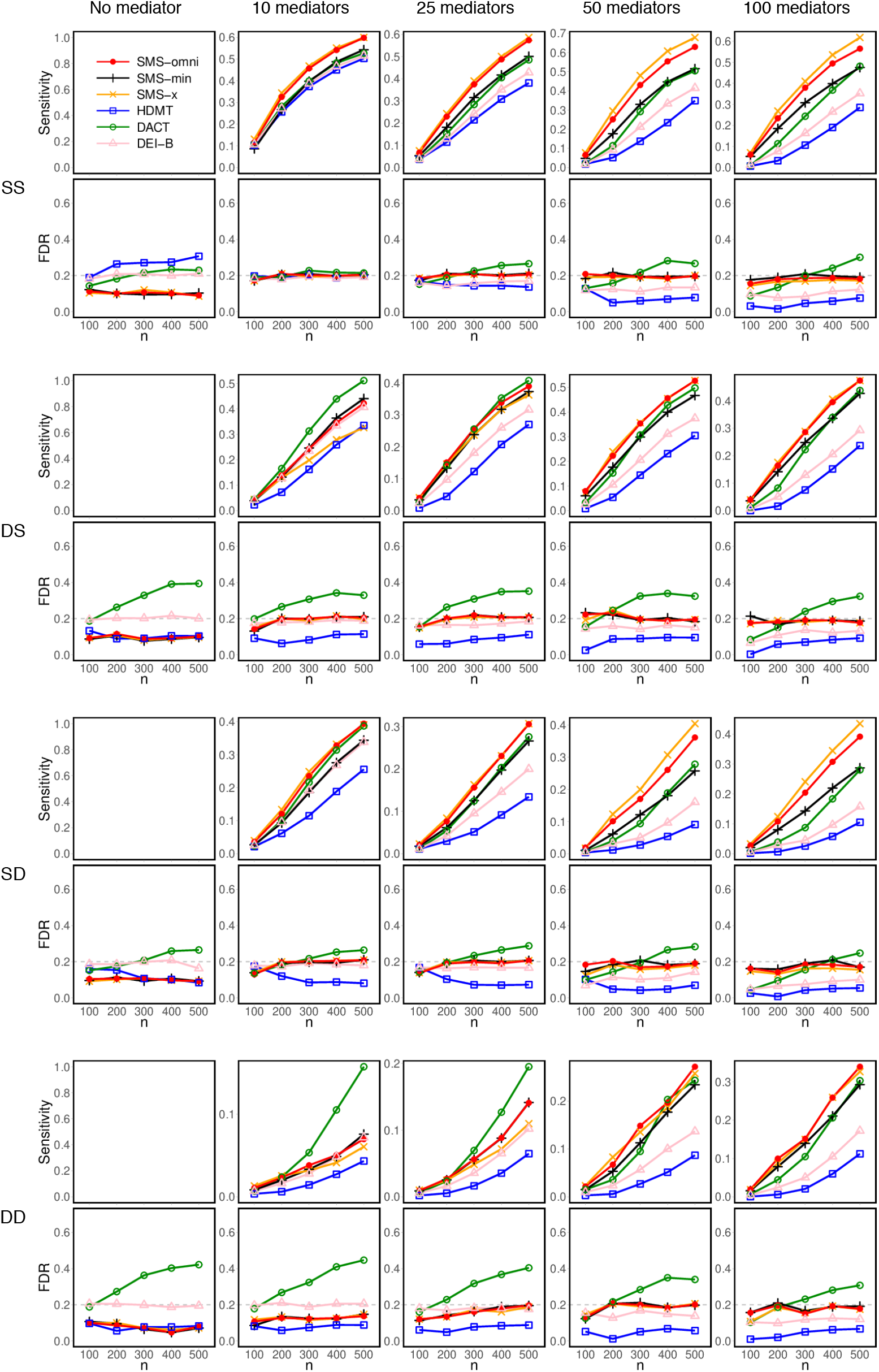
Simulation results on empirical FDR and sensitivity, based on *J* = 1,000 features. The dashed gray line indicates the nominal FDR level of 0.2.

**Figure 5:**
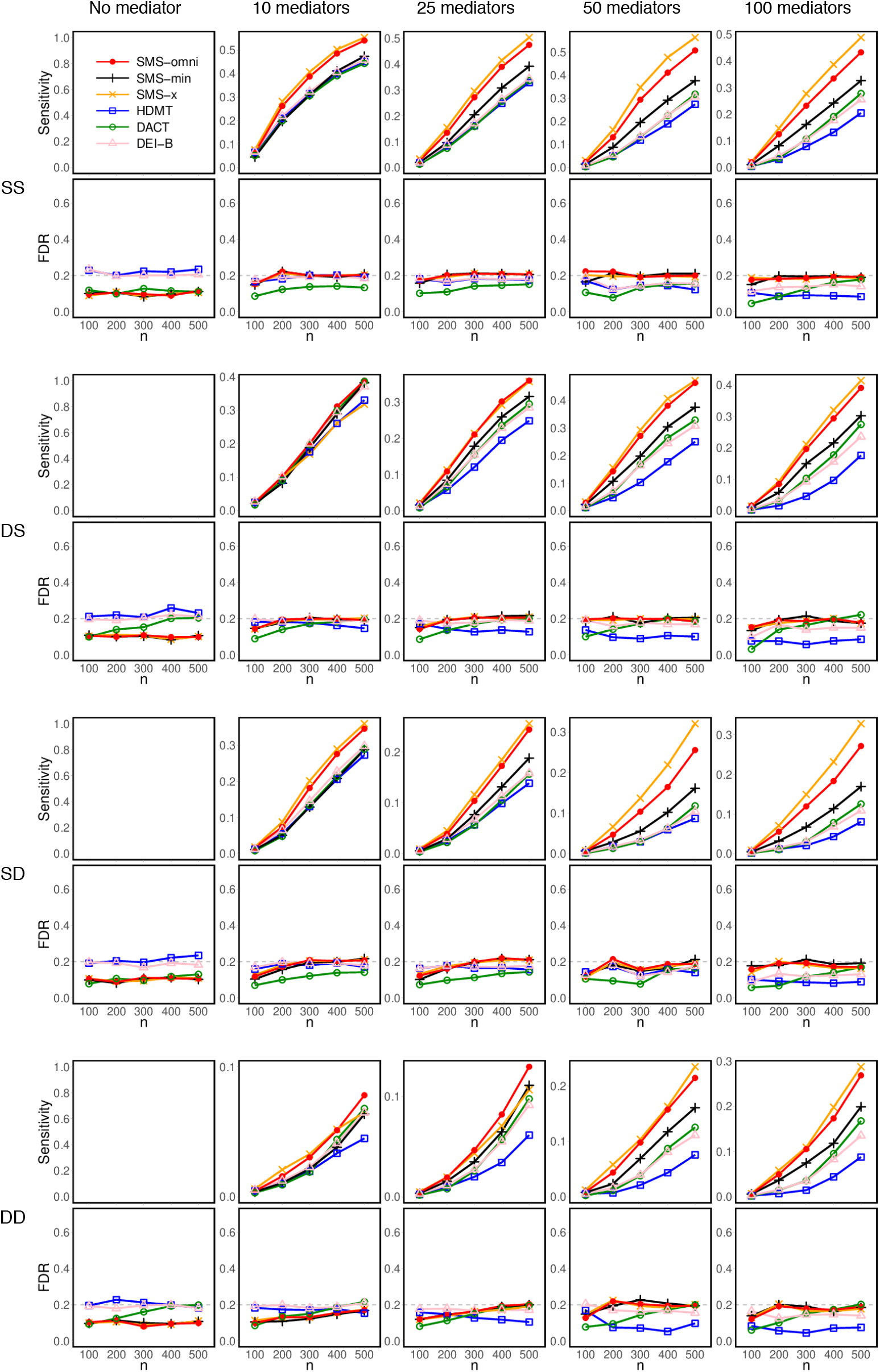
Simulation results on empirical FDR and sensitivity, based on *J* = 5,000 features. The dashed gray line indicates the nominal FDR level of 0.2.

**Figure 6:**
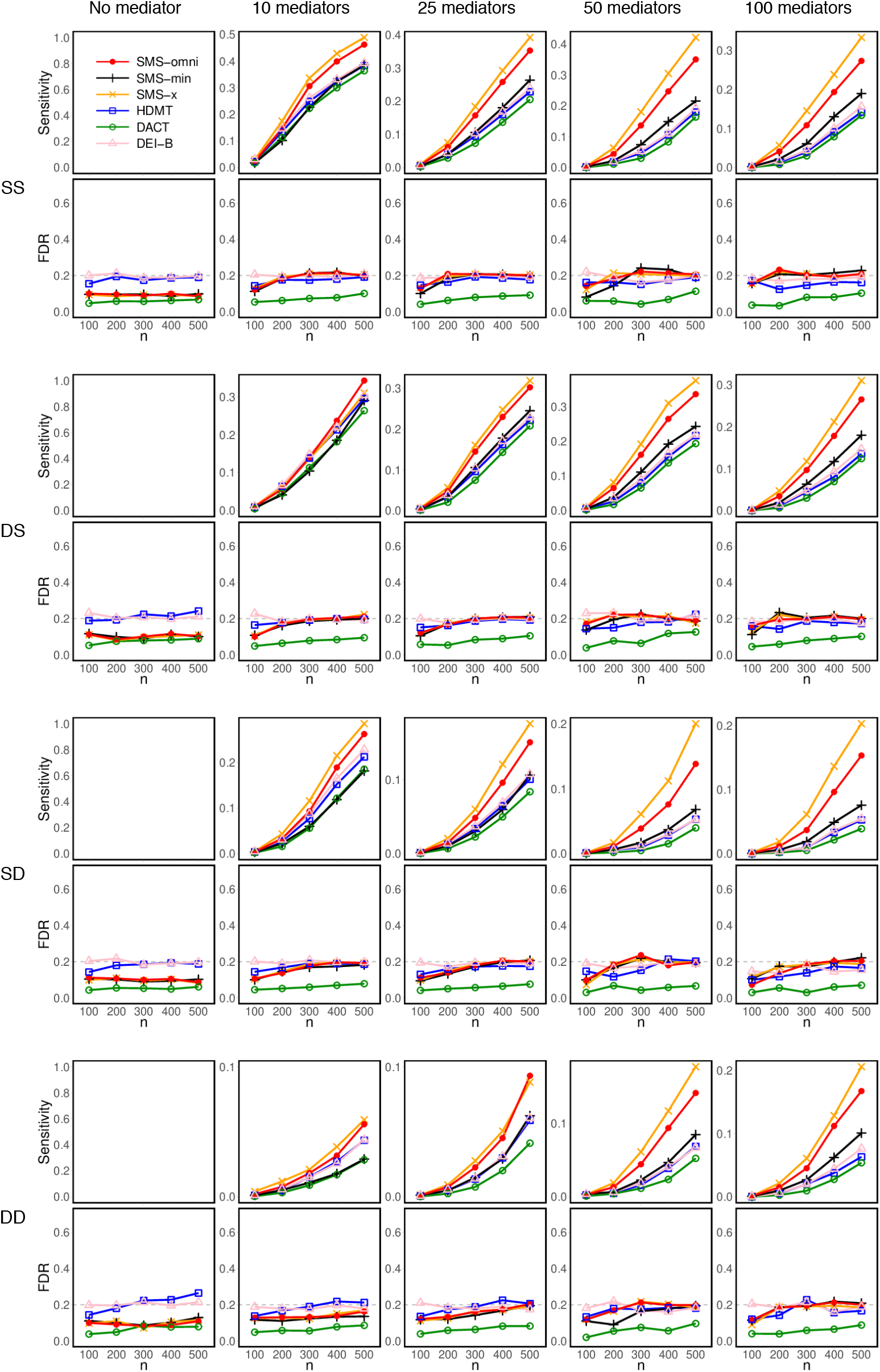
Simulation results on empirical FDR and sensitivity, based on J = 50,000 features. The dashed gray line indicates the nominal FDR level of 0.2.

The SMS methods generally achieved the highest sensitivity among methods that controlled the FDR. The sensitivity gain was especially pronounced in dense-mediator settings, with 50 or 100 true mediators. For example, when SMS-omni achieved adequate sensitivity, such as 50%, its sensitivity gain over DEI-B often exceeded 20 percentage points. In sparse-mediator settings with only 10 true mediators, the sensitivity gain was small, especially under the DS and DD settings, although SMS-omni remained at least as sensitive as DEI-B. Because SMS-min uses the same statistics as DEI-B, the sensitivity gain of SMS-min over DEI-B reflects improved FDR calibration. We selected DEI-B as the primary benchmark because it consistently outperformed HDMT and DACT, achieving the highest sensitivity among existing methods while maintaining FDR control across settings.

Among the SMS methods, as expected, SMS-omni consistently tracked the better sensitivity between SMS-x and SMS-min. In most settings, especially with dense mediators, SMS-x yielded substantially higher sensitivity than SMS-min. In a few cases where mediators were sparse, type-I or type-II nulls were dense, and *J* was relatively small, SMS-min was more sensitive than SMS-x.

Figure 7 presents results under varying relative strengths of the E–M and M–O associations. We considered two representative scenarios that resemble the configurations observed in the real data: a lower-dimensional setting with *J* = 1,000 under SD, and a high-dimensional setting with *J* = 50,000 under DS. As the association strengths became increasingly imbalanced, SMS-x progressively outperformed SMS-min.

**Figure 7:**
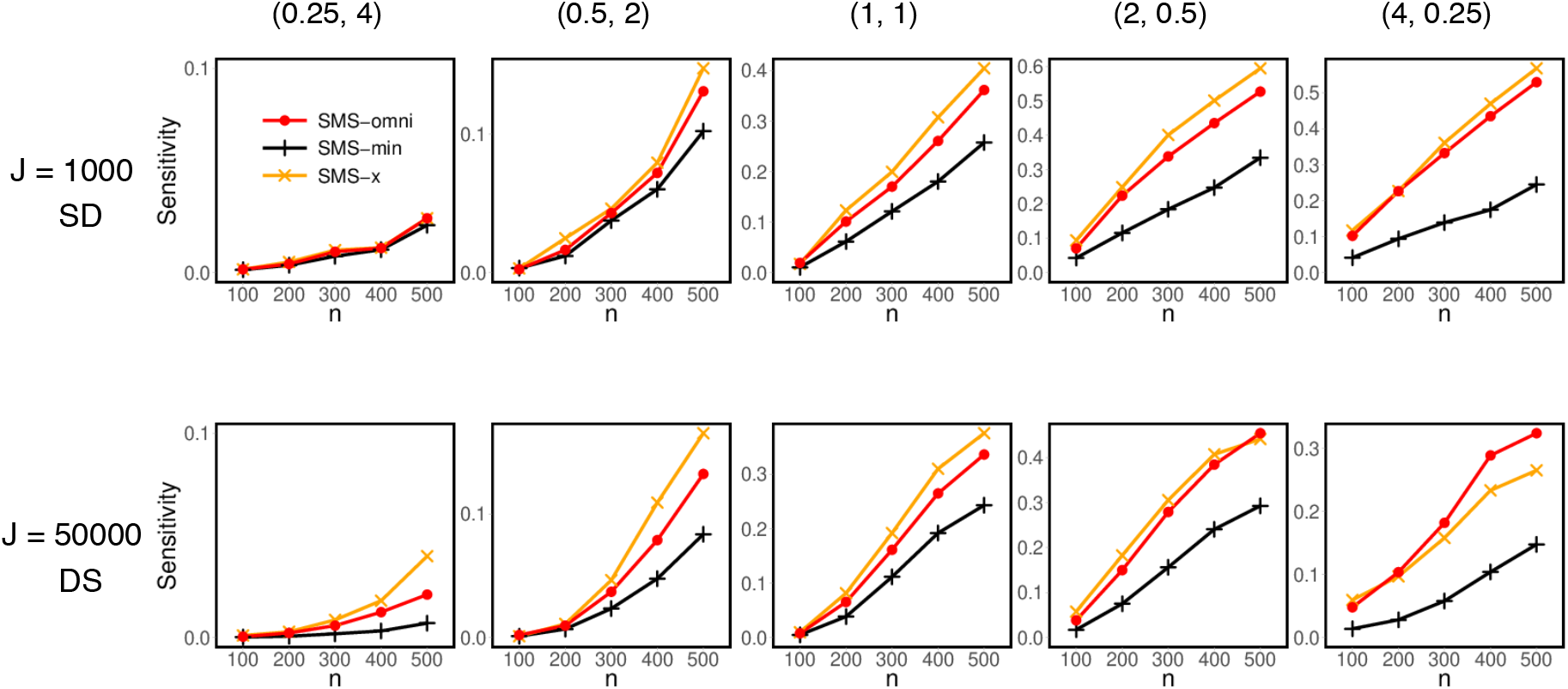
Simulation results (with 50 true mediators) on the sensitivity of the SMS methods under varying relative strengths of the E–M and M–O associations, governed by different values of (*c*_1_, *c*_2_), across two settings that mimic the real datasets.

Figure S5 shows that using 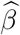-based statistics led to only a slight loss of sensitivity relative to using *Z*-scores; the difference was particularly small for SMS-x and, consequently, for SMS-omni. Figures S6 and S7 further show that the results for a single binary outcome and feature-specific continuous outcomes followed patterns similar to those observed for a single continuous outcome (Figure 4).

### PKU-SPCO metabolomics study

Returning to the PKU-SPCO metabolomics study of childhood obesity, we analyzed whether and how maternal obesity, a binary exposure, influences childhood obesity, a binary outcome, through alterations in metabolite abundances. As described earlier, the dataset comprises 128 children, including 70 with obesity and 58 with normal weight, with abundance measurements for 1,152 metabolites, along with parental, demographic, lifestyle, and clinical covariates. The data first confirmed a positive association between maternal obesity and childhood obesity, with an odds ratio of 5.5 and a chi-squared *p*-value of 2.1 × 10^−5^. Because the total effect was positive, the outcome coding was retained without reversal.

Prior to analysis, the metabolomics data were preprocessed by replacing zero values with one-half of the smallest observed positive value for each metabolite, followed by a centered log-ratio (CLR) transformation [27] to account for compositionality. For each metabolite, we fitted a linear regression model of metabolite abundance on maternal obesity and a logistic regression model of childhood obesity on metabolite abundance and maternal obesity. Both models were adjusted for household income and daily total caloric intake. Household income was dichotomized at CNY 200,000 per year, and caloric intake was treated as continuous.

Figure 2 shows the distributions of the *p*-values for the E–M and M–O associations. The stronger deviation from the uniform reference among the M–O *p*-values suggests that the children’s metabolome was more strongly associated with childhood obesity than with maternal obesity. Figure 3 shows the distributions of the SMS statistics, all of which exhibit a heavier positive tail, suggesting the presence of plausible mediators.

We applied SMS-omni to the *Z*-scores and HDMT, DEI-B, and DACT to the *p*-values to identify mediators at a nominal FDR level of 0.2. As shown in Table 2, none of the existing methods detected significant mediators. Indeed, applying the Benjamini–Hochberg (BH) procedure [28] at FDR level of 0.2 to either set of marginal *p*-values identified no significant E–M or M–O associations (Table 2). In contrast, SMS-omni identified five significant mediators, with details provided in Table 3. These mediators included not only metabolites with small *p*-values for both associations, but also one metabolite, ranked third, for which one marginal *p*-value was only near significant (0.0535) while the other was highly significant (0.0021). This example illustrates how SMS can detect mediators supported by the joint evidence of two associations, even when one marginal association alone would not pass conventional significance thresholds.

**Table 2:** Number of mediators identified in the real datasets.

| Dataset | $n$ | $J$ | Identified<br>E–M<br>associations* | Identified<br>M–O<br>associations* | Method | Identified<br>mediators** |
| --- | --- | --- | --- | --- | --- | --- |
| PKU-SPCO<br>metabolomics<br>data | 128 | 1,152 | 0 | 0 | SMS-omni | 5 |
|  |  |  |  |  | HDMT | 0 |
|  |  |  |  |  | DEI-B | 0 |
|  |  |  |  |  | DACT | 0 |
| NAS<br>DNA methylation<br>data | 603 | 484,613 | 36 | 10 | SMS-omni | 70 |
|  |  |  |  |  | HDMT | 25 |
|  |  |  |  |  | DEI-B | 24 |
|  |  |  |  |  | DACT | 19 |

Note: \* Identified by applying the BH procedure to the $p$ -values at a nominal FDR level of 0.2. \*\* Obtained at FDR level 0.2 for the PKU-SPCO dataset and 0.05 for the NAS dataset. The mediator sets identified by different methods in the NAS data exhibit a strictly nested structure as described in the main text.

**Table 3:**
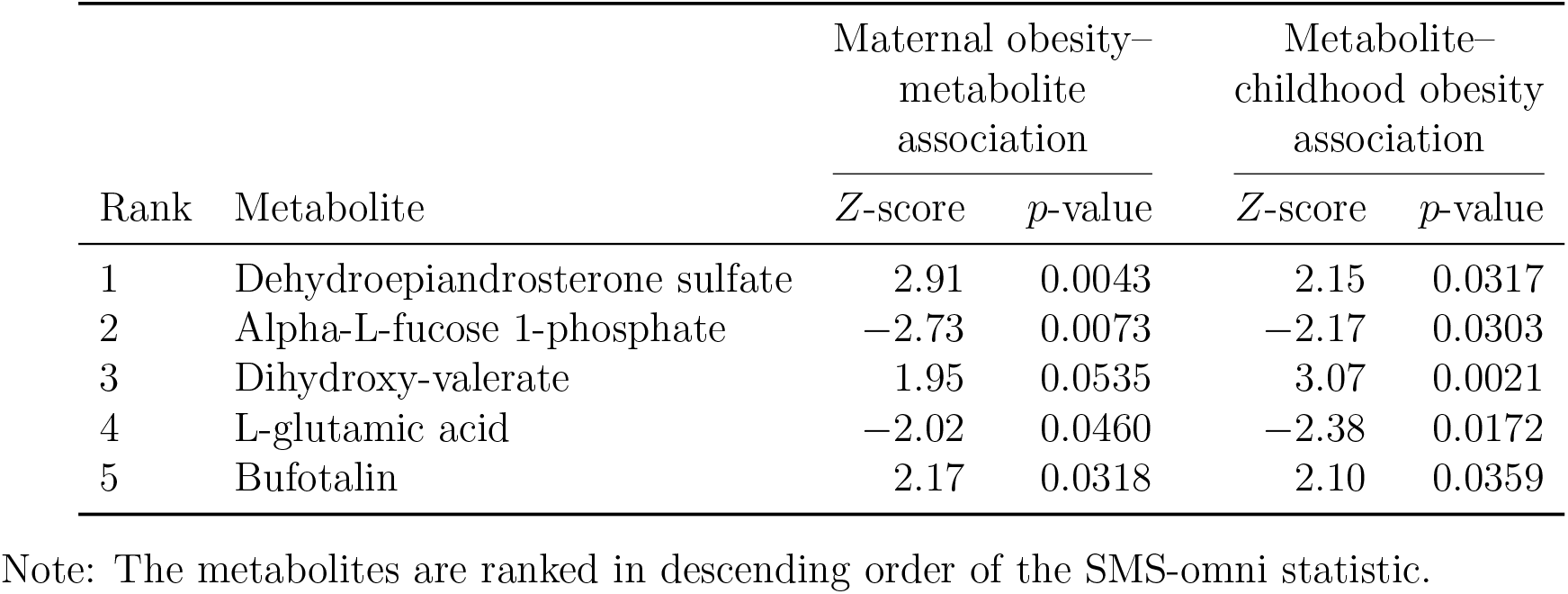
Metabolites identified by SMS-omni in the PKU-SPCO dataset.

| Rank | Metabolite | Maternal obesity–<br>metabolite<br>association |  | Metabolite–<br>childhood obesity<br>association |  |
| --- | --- | --- | --- | --- | --- |
|  |  | *Z*-score | *p*-value | *Z*-score | *p*-value |
| 1 | Dehydroepiandrosterone sulfate | 2.91 | 0.0043 | 2.15 | 0.0317 |
| 2 | Alpha-L-fucose 1-phosphate | −2.73 | 0.0073 | −2.17 | 0.0303 |
| 3 | Dihydroxy-valerate | 1.95 | 0.0535 | 3.07 | 0.0021 |
| 4 | L-glutamic acid | −2.02 | 0.0460 | −2.38 | 0.0172 |
| 5 | Bufotalin | 2.17 | 0.0318 | 2.10 | 0.0359 |

Note: The metabolites are ranked in descending order of the SMS-omni statistic.

Previous studies suggested that maternal obesity may shape early offspring gut metabolic profiles through intestinal colonization, fecal short-chain fatty acids, and human milk oligosaccharide (HMO)-microbiome interactions, thereby contributing to childhood adiposity risk [29– 31]. The five metabolites identified here point to several plausible mediating pathways. Dehydroepiandrosterone sulfate (DHEAS) is a host steroid precursor, and may play a mediating role consistent with prior evidence linking childhood DHEAS with adiposity, insulin, and cardiometabolic traits [32]. Alpha-L-fucose 1-phosphate may reflect fucose utilization and fucosylated oligosaccharide-related metabolism, a pathway plausibly affected by maternal obesity through altered HMO composition and infant gut development [30, 31, 33]. Dihydroxyvalerate may represent hydroxy fatty acid–related metabolism associated with microbial fermentation, fatty acid oxidation, or broader intestinal metabolic activity [34, 35]. L-glutamic acid is involved in amino acid metabolism and has been associated with insulin resistance in pediatric obesity [36]. Bufotalin may be related to oxidative-stress pathways, as experimental studies suggest effects on GPX4 degradation, lipid peroxidation, and ferroptosis [37, 38]. Overall, these findings provide hypothesis-generating evidence that steroid metabolism, organic acid metabolism, amino acid metabolism, fucose utilization, and oxidative-stress-related gut metabolic pathways may mediate the association between maternal obesity and childhood obesity risk.

### Normative Aging DNA methylation study

These data were previously analyzed by DACT and DEI-B, enabling direct comparison between these methods and the SMS methods. The data were obtained from the Normative Aging Study [23] and include DNA methylation measurements for 484,613 CpG sites from 603 individuals. The goal was to investigate whether the effect of smoking, a binary exposure, on lung function, a continuous outcome, is mediated through DNA methylation at specific CpG sites. Methylation levels at each CpG site were originally measured as Beta values and then transformed into M-values using the logit function with base 2.

We obtained summary statistics, including effect-size estimates, *Z*-scores, and *p*-values, for the association between smoking and DNA methylation at each CpG site, and for the association between CpG methylation and lung function conditional on smoking. Both associations were adjusted for age, height, weight, education history, medication history, blood cell-type abundances, and five principal components. Because smoking was inversely associated with lung function, we flipped the signs of the *Z*-scores for the CpG methylation–lung function associations.

Figure 2 shows that the *p*-values for the E–M associations are enriched near zero, suggesting substantial effects of smoking on DNA methylation at multiple CpG sites. In contrast, the *p*-values for the M–O associations are approximately uniform, or even sub-uniform, indicating weaker associations between DNA methylation and lung function. Consistently, applying the BH procedure at the nominal FDR level of 0.2 identified 36 significant E–M associations and 10 M–O associations (Table 2). The distributions of the SMS statistics are presented in Figure 3. Owing to the much larger number of features in this dataset, all three distributions exhibit a highly symmetric component centered around zero, corresponding to the composite nulls. In addition, all three display a pronounced positive tail, suggesting the presence of plausible mediators.

We applied SMS-omni to the *Z*-scores and HDMT, DEI-B, and DACT to the *p*-values to identify mediators. The nominal FDR level was set to 0.05 for consistency with previous studies. As shown in Table 2, SMS-omni identified 70 mediators. In comparison, HDMT, DEI-B, and DACT identified 25, 24, and 19 mediators, respectively, with the latter two matching the findings reported in the original DEI-B and DACT papers. Notably, all CpG sites identified by the existing methods were also identified by SMS-omni. Moreover, the detected mediator sets exhibit a strict nested structure: the set identified by SMS-omni contained that identified by HDMT, which contained that identified by DEI-B, which in turn contained that identified by DACT. These patterns align with the simulation results for *J* = 50,000, where SMS-omni achieved the highest sensitivity, followed by HDMT and DEI-B with similar sensitivity, while DACT showed the lowest sensitivity.

Detailed information on the identified mediators is summarized in Table S1. Notably, among the mediators identified by the existing methods, all corresponding products *Z*_1*j*_*Z*_2*j*_ had the same sign, even though this is not required by those methods. This pattern supports the sign-consistency assumption underlying the SMS methods. The additional 45 mediators identified by SMS-omni, but not by any existing methods, typically exhibited one moderately significant *p*-value together with another highly significant *p*-value, highlighting the advantage of SMS-x.

## Discussions

We proposed SMS, a new framework for high-dimensional mediation analysis that exploits the symmetry of mediation statistics to calibrate the composite null distribution as a whole. This approach differs fundamentally from traditional methods that explicitly model the composite-null distribution. A key advantage of SMS is its improved power, particularly when the strengths of the E–M and M–O associations are imbalanced. The SMS framework allows flexible combination of evidence from the two associations, and enables construction of an omnibus test for practical use. This property is especially relevant for mediation analysis, where the two associations are often imbalanced, unlike in data-splitting settings where the two components are typically more symmetric.

The symmetric-statistic framework works best when both the number of features and the number of true signals are large. To evaluate its performance under more challenging and practically relevant conditions, our simulations included settings with as few as 1,000 features, comparable to the PKU-SPCO metabolomics dataset, and as few as 10 true mediators. Although we described settings with 50 or more true mediators as dense, this designation is conservative relative to prior studies. For instance, HDMT defined 20% true mediators as dense and 2% as sparse, which correspond to 1,000 and 100 mediators, respectively, even in its lowest-dimensional setting of 5,000 features. Similarly, DACT considered 41 true mediators, and the sparsest-signal setting examined in DEI-B included 50 true mediators. Thus, our simulation design placed particular emphasis on sparse-signal scenarios.

High-dimensional features, including metabolites, gene expression levels, and microbes, often exhibit complex feature–feature dependence. Methods such as DACT and DEI-B rely on the BH procedure for multiple-testing correction, which requires independence or certain forms of positive dependence among features [39]. In contrast, the symmetric-statistic framework can accommodate more flexible dependence structures under a weak-dependence condition [22]. Moreover, empirical evidence from Dai et al. [22] suggests that the framework can maintain satisfactory FDR control even when this condition is not strictly satisfied.

The proposed SMS methods were implemented in the R package, SMStat, available on CRAN and GitHub at https://github.com/yijuanhu/SMStat. The implementation is highly efficient. As shown in Figure S8, although DEI-B was the fastest method, the SMS methods completed both real-data analyses in fractions of a second and were substantially faster than HDMT and DACT. Computation times were measured after the effect-size estimates, *Z*-scores, and *p*-values for the two associations had been obtained. All timings were based on runs performed on a MacBook Air with an Apple M3 chip and 16 GB memory.

Our methods use the null statistics to construct a background distribution against which statistics under the alternative are contrasted. Therefore, a larger number of null features provides a more stable estimate of this background distribution. In this sense, high dimensionality becomes an advantage rather than a burden. For this reason, we do not recommend pre-screening procedures that remove null features before analysis [9]. Nevertheless, our methods remain applicable to moderately sized datasets; for example, they performed well with as few as *J* = 500 features (Figure S9).

## References

1. Yan S, Liu Y, Yin L, Zhang L, Duan X, Liang B, et al. Factors influencing preschool children’s obesity in two cities of Shandong Province. Chinese Journal of Child Health Care. 2025;33(2):142–148.

2. Godfrey KM, Reynolds RM, Prescott SL, Nyirenda M, Jaddoe VWV, Eriksson JG, et al. Influence of maternal obesity on the long-term health of offspring. The Lancet Diabetes & Endocrinology. 2017;5(1):53–64.

3. Fleming TP, Watkins AJ, Velazquez MA, Mathers JC, Prentice AM, Stephenson J, et al. Origins of lifetime health around the time of conception: causes and consequences. Lancet. 2018;391(10132):1842–1852.

4. Creanga AA, Catalano PM, Bateman BT. Obesity in Pregnancy. New England Journal of Medicine. 2022;387(3):248–259.

5. Sobel ME. Asymptotic confidence intervals for indirect effects in structural equation models. Sociological methodology. 1982;13:290–312.

6. MacKinnon DP, Lockwood CM, Hoffman JM, West SG, Sheets V. A comparison of methods to test mediation and other intervening variable effects. Psychological methods. 2002;7(1):83.

7. Dai JY, Stanford JL, LeBlanc M. A multiple-testing procedure for high-dimensional mediation hypotheses. Journal of the American Statistical Association. 2022;117(537):198– 213.

8. Storey JD, Taylor JE, Siegmund D. Strong Control, Conservative Point Estimation and Simultaneous Conservative Consistency of False Discovery Rates: A Unified Approach. Journal of the Royal Statistical Society Series B: Statistical Methodology. 2004 02;66(1):187–205.

9. Liu Z, Shen J, Barfield R, Schwartz J, Baccarelli AA, Lin X. Large-scale hypothesis testing for causal mediation effects with applications in genome-wide epigenetic studies. Journal of the American Statistical Association. 2022;117(537):67–81.

10. Jin J, Cai TT. Estimating the Null and the Proportion of Nonnull Effects in Large-Scale Multiple Comparisons. Journal of the American Statistical Association. 2007;102(478):495–506.

11. Efron B, Tibshirani R, Storey JD, Tusher V. Empirical Bayes Analysis of a Microarray Experiment. Journal of the American Statistical Association. 2001;96(456):1151–1160.

12. Du J, Zhou X, Clark-Boucher D, Hao W, Liu Y, Smith JA, et al. Methods for large-scale single mediator hypothesis testing: possible choices and comparisons. Genetic epidemiology. 2023;47(2):167–184.

13. Liu Y. A simple and powerful method for large-scale composite null hypothesis testing with applications in mediation analysis. Biometrics. 2025;81(1):ujaf011.

14. Stratakis N, Anguita-Ruiz A, Fabbri L, Maitre L, González JR, Andrusaityte S, et al. Multi-omics architecture of childhood obesity and metabolic dysfunction uncovers biological pathways and prenatal determinants. Nature Communications. 2025;16(1):654.

15. Santos Ferreira DL, Williams DM, Kangas AJ, Soininen P, Ala-Korpela M, Smith GD, et al. Association of pre-pregnancy body mass index with offspring metabolic profile: Analyses of 3 European prospective birth cohorts. PLOS Medicine. 2017 08;14(8):1–19.

16. Lowe J William L, Bain JR, Nodzenski M, Reisetter AC, Muehlbauer MJ, Stevens RD, et al. Maternal BMI and Glycemia Impact the Fetal Metabolome. Diabetes Care. 2017 06;40(7):902–910.

17. Catalano PM, Shankar K. Obesity and pregnancy: mechanisms of short term and long term adverse consequences for mother and child. BMJ. 2017;356.

18. Hu YJ, Satten GA. Testing hypotheses about the microbiome using the linear decomposition model (LDM). Bioinformatics. 2020;36(14):4106–4115.

19. Hu Y, Satten GA, Hu YJ. LOCOM: A logistic regression model for testing differential abundance in compositional microbiome data with false discovery rate control. Proceedings of the National Academy of Sciences. 2022 7;119(30).

20. Barber RF, Candès EJ. Controlling the false discovery rate via knockoffs. The Annals of Statistics. 2015;43(5):2055 – 2085.

21. Candès E, Fan Y, Janson L, Lv J. Panning for Gold: ‘Model-X’ Knockoffs for High Dimensional Controlled Variable Selection. Journal of the Royal Statistical Society Series B: Statistical Methodology. 2018 06;80(3):551–577.

22. Dai C, Lin B, Xing X, Liu JS. False discovery rate control via data splitting. Journal of the American Statistical Association. 2023;118(544):2503–2520.

23. Bell B, Rose CL, Damon A. The Normative Aging Study: An Interdisciplinary and Longitudinal Study of Health and Aging. The International Journal of Aging and Human Development. 1972;3(1):5–17.

24. VanderWeele T, Vansteelandt S. Conceptual issues concerning mediation, interventions and composition. Statistics and Its Interface. 2009;2:457–468.

25. VanderWeele T, Vansteelandt S. Mediation analysis with multiple mediators. Epidemiologic methods. 2014;2(1):95–115. PMCID: PMC4287269.

26. MacKinnon D, Dwyer J. Estimating Mediated Effects in Prevention Studies. Evaluation Review. 1993;17:144–158.

27. Aitchison J. The statistical analysis of compositional data. Chapman and Hall, London-New York; 1986.

28. Benjamini Y, Hochberg Y. Controlling the false discovery rate: a practical and powerful approach to multiple testing. Journal of the royal statistical society Series B (Methodological). 1995;p. 289–300.

29. Gilley SP, Ruebel ML, Sims C, Zhong Y, Turner D, Lan RS, et al. Associations between maternal obesity and offspring gut microbiome in the first year of life. Pediatric Obesity. 2022;17(9):e12921.

30. Saben JL, Sims CR, Abraham A, Bode L, Andres A. Human Milk Oligosaccharide Concentrations and Infant Intakes Are Associated with Maternal Overweight and Obesity and Predict Infant Growth. Nutrients. 2021;13(2).

31. Corona-Cervantes K, Urrutia-Baca VH, Gámez-Valdez JS, Jiménez-López B, Rodríguez-Gutierrez NA, Chávez-Caraza K, et al. Maternal obesity alters human milk oligosaccharides content and correlates with early acquisition of late colonizers in the neonatal gut microbiome. Gut Microbes. 2026;18(1):2607043.

32. Santos-Silva R, Fontoura M, Guimarães JT, Barros H, Santos AC. Association of dehydroepiandrosterone sulfate, birth size, adiposity and cardiometabolic risk factors in 7-year-old children. Pediatr Res. 2022;91(7):1897–1905.

33. Korpela K, Salonen A, Hickman B, Kunz C, Sprenger N, Kukkonen K, et al. Fucosylated oligosaccharides in mother’s milk alleviate the effects of caesarean birth on infant gut microbiota. Scientific Reports. 2018;8(1):13757.

34. Zhang S, Dang Y. Roles of gut microbiota and metabolites in overweight and obesity of children. Front Endocrinol (Lausanne). 2022;13:994930.

35. Mann ER, Lam YK, Uhlig HH. Short-chain fatty acids: linking diet, the microbiome and immunity. Nature Reviews Immunology. 2024;24(8):577–595.

36. Gumus Balikcioglu P, Jachthuber Trub C, Balikcioglu M, Ilkayeva O, White PJ, Muehlbauer M, et al. Branched-chain α-keto acids and glutamate/glutamine: Biomarkers of insulin resistance in childhood obesity. Endocrinol Diabetes Metab. 2023;6(1):e388.

37. Zhang W, Jiang B, Liu Y, Xu L, Wan M. Bufotalin induces ferroptosis in non-small cell lung cancer cells by facilitating the ubiquitination and degradation of GPX4. Free Radical Biology and Medicine. 2022;180:75–84.

38. Ursini F, Maiorino M. Lipid peroxidation and ferroptosis: The role of GSH and GPx4. Free Radic Biol Med. 2020;152:175–185.

39. Benjamini Y, Yekutieli D. The Control of the False Discovery Rate in Multiple Testing under Dependency. The Annals of Statistics. 2001;29(4):1165–1188.

