## Supplemental materials for "SMS: Symmetric Mediation Statistics for Powerful High-Dimensional Mediation Analysis"

### Supplementary Materials

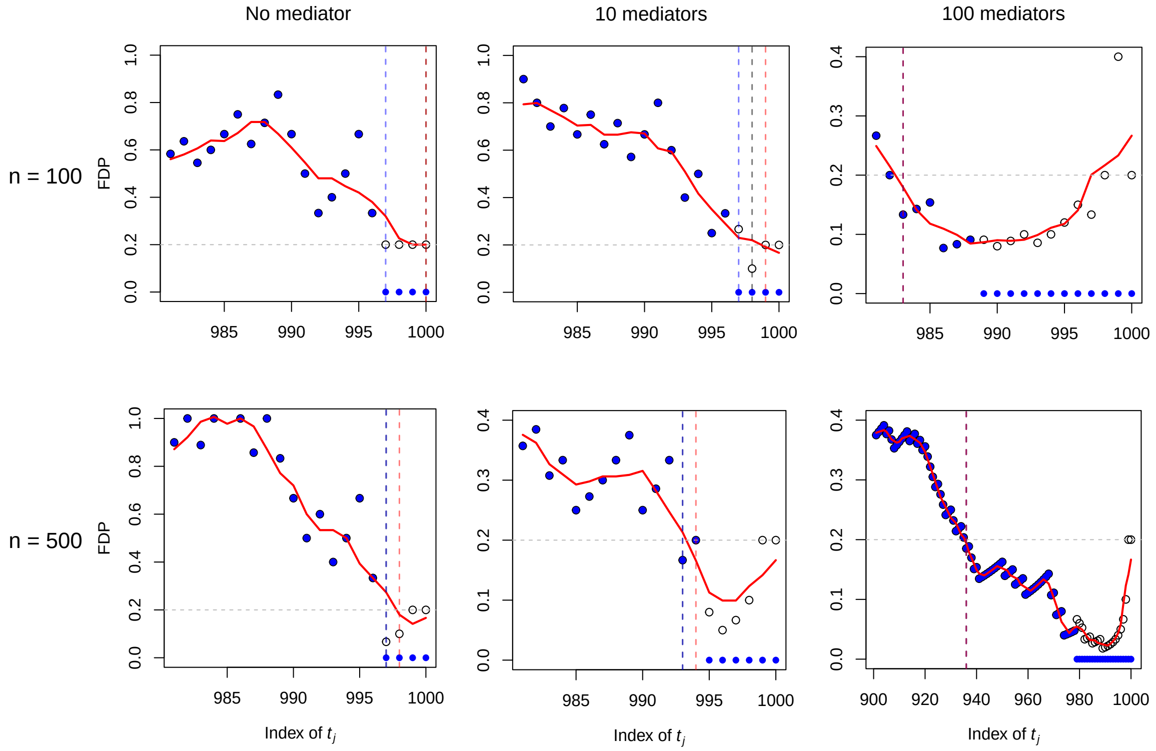


Figure S1: Comparison of the three FDP estimators for the SMS-x method: $\widehat{\text{FDP}}(t)$ (blue solid points), $\widetilde{\text{FDP}}(t)$ (black circles), and $\widetilde{\text{FDP}}_W(t)$ (the red line). Each vertical dashed line indicates the threshold $\tau_\alpha$ selected by the FDP estimator of the same color. The gray dotted line denotes the nominal FDR level, $\alpha = 0.2$. The $t_j$'s are the ordered values of $\{|S_1|, |S_2|, \ldots, |S_J|\}$. Each plot is based on one simulated dataset with $J = 1000$ under the SD setting.

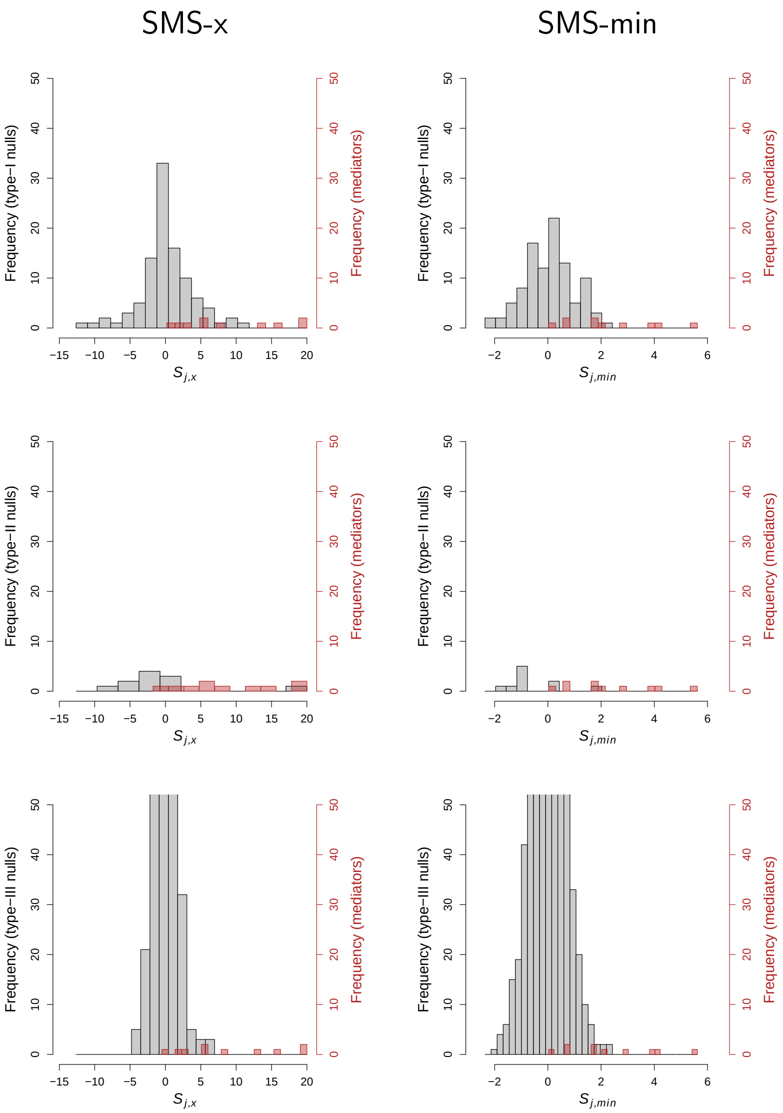


Figure S2: Distributions of the SMS-x and SMS-min statistics for features under the three null types and for true mediators, based on a simulated dataset with $n = 500$, $J = 1{,}000$, and 10 mediators under the DS setting. The red region denotes the true mediators. The x-axis for SMS-x is truncated at 20, and the y-axis for the type-III null panel is truncated at 50, to show greater detail in the positive tail of the gray region.

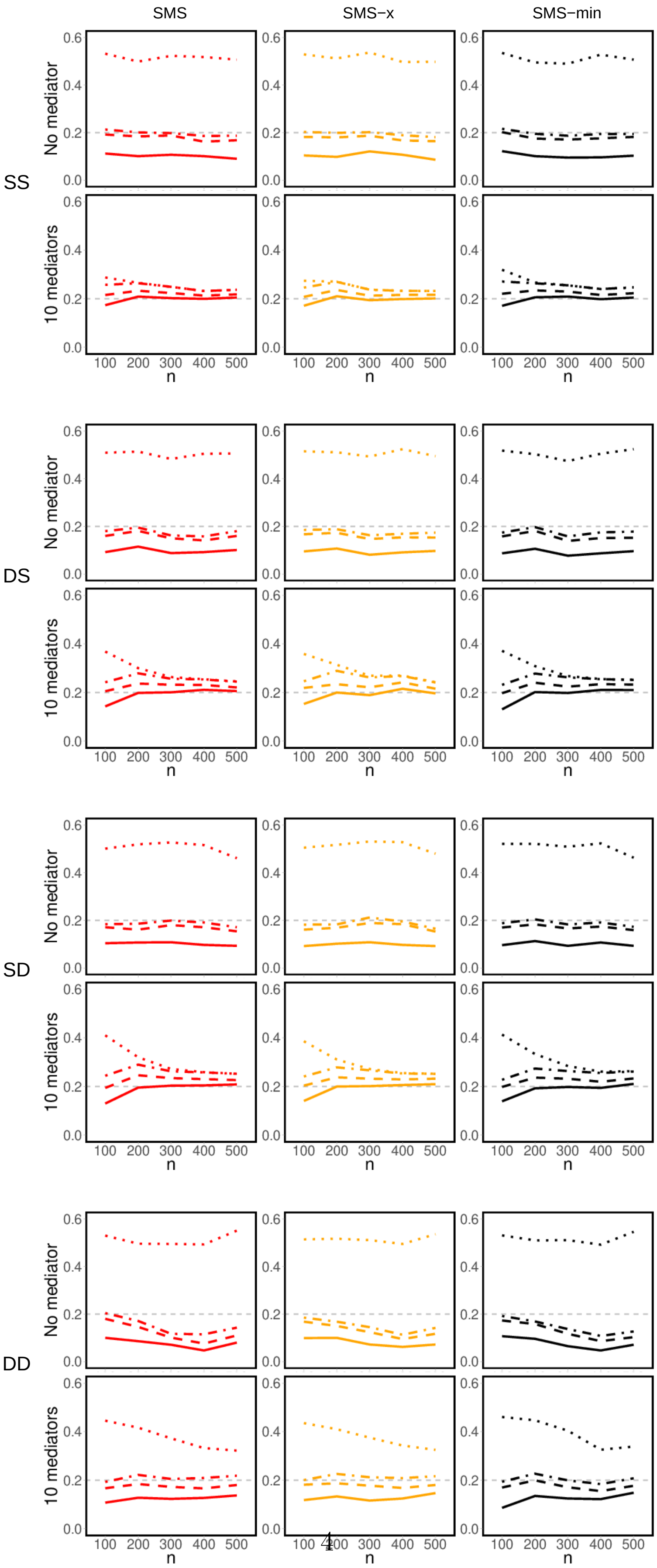




Figure S3: Simulation results ($J = 1{,}000$) on empirical FDR for different variants of the SMS methods: no correction (dotted line), correction without stabilization (dash-dotted

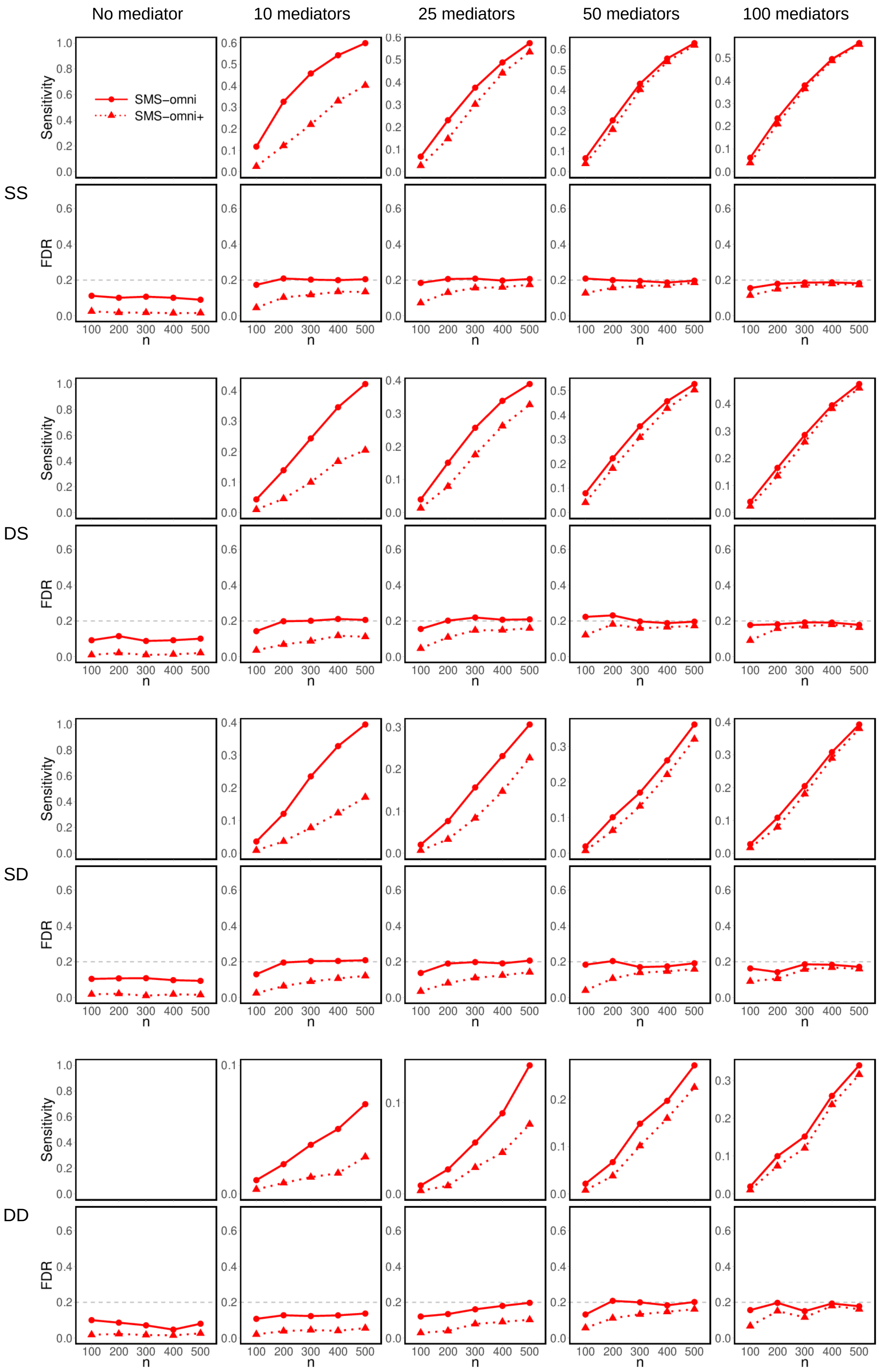




Figure S4: Simulation results ($J = 1{,}000$) comparing SMS-omni, which uses the proposed FDP estimator in (8), with SMS-omni+, which uses the conservative FDP estimator in (9). The nominal FDR level of 0.2 is indicated by a dashed gray line.

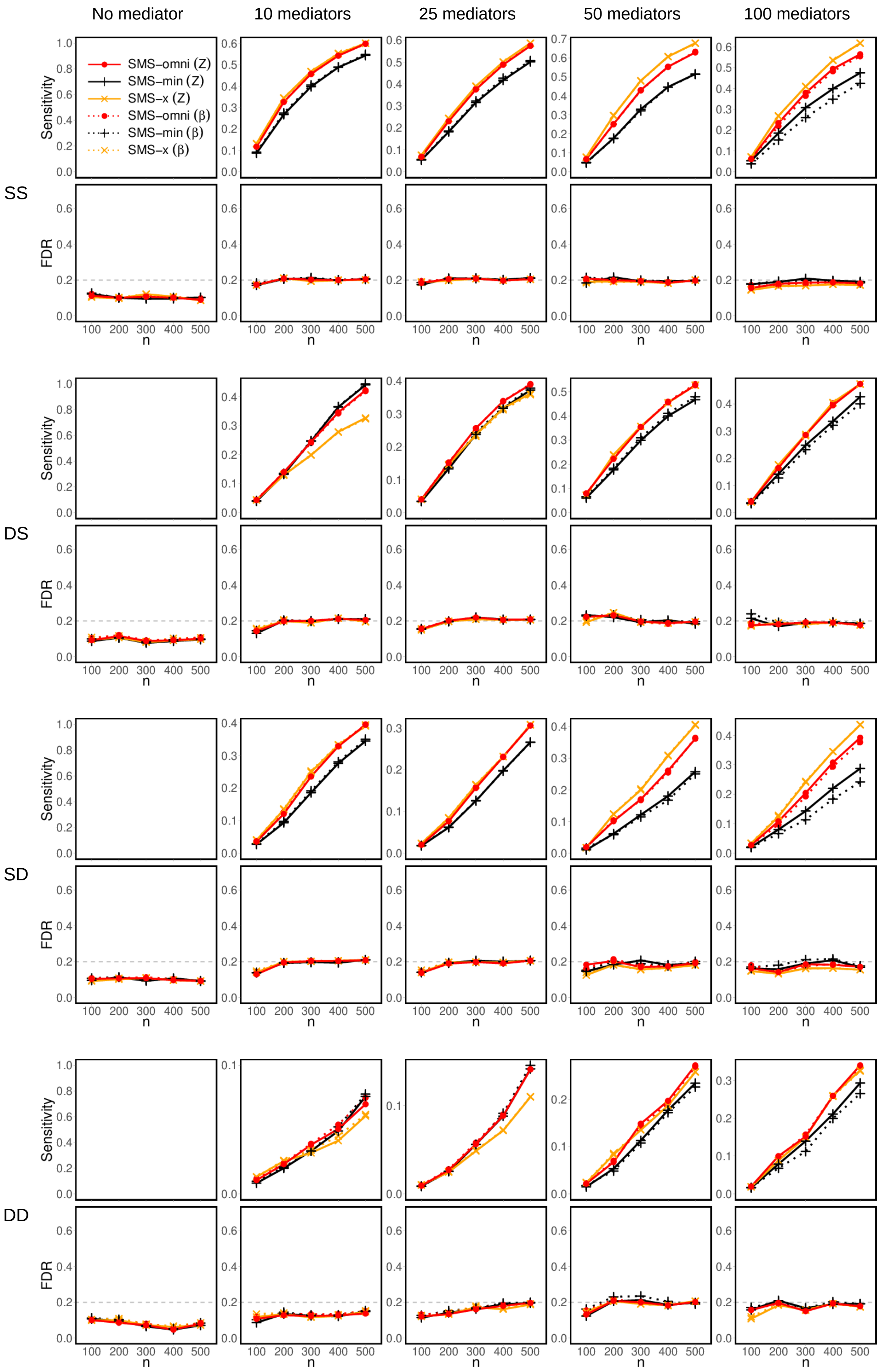




Figure S5: Simulation results ($J = 1{,}000$) comparing $\widehat{\beta}$-based statistics with $Z$-score-based statistics. The dashed gray line indicates the nominal FDR level of 0.2.

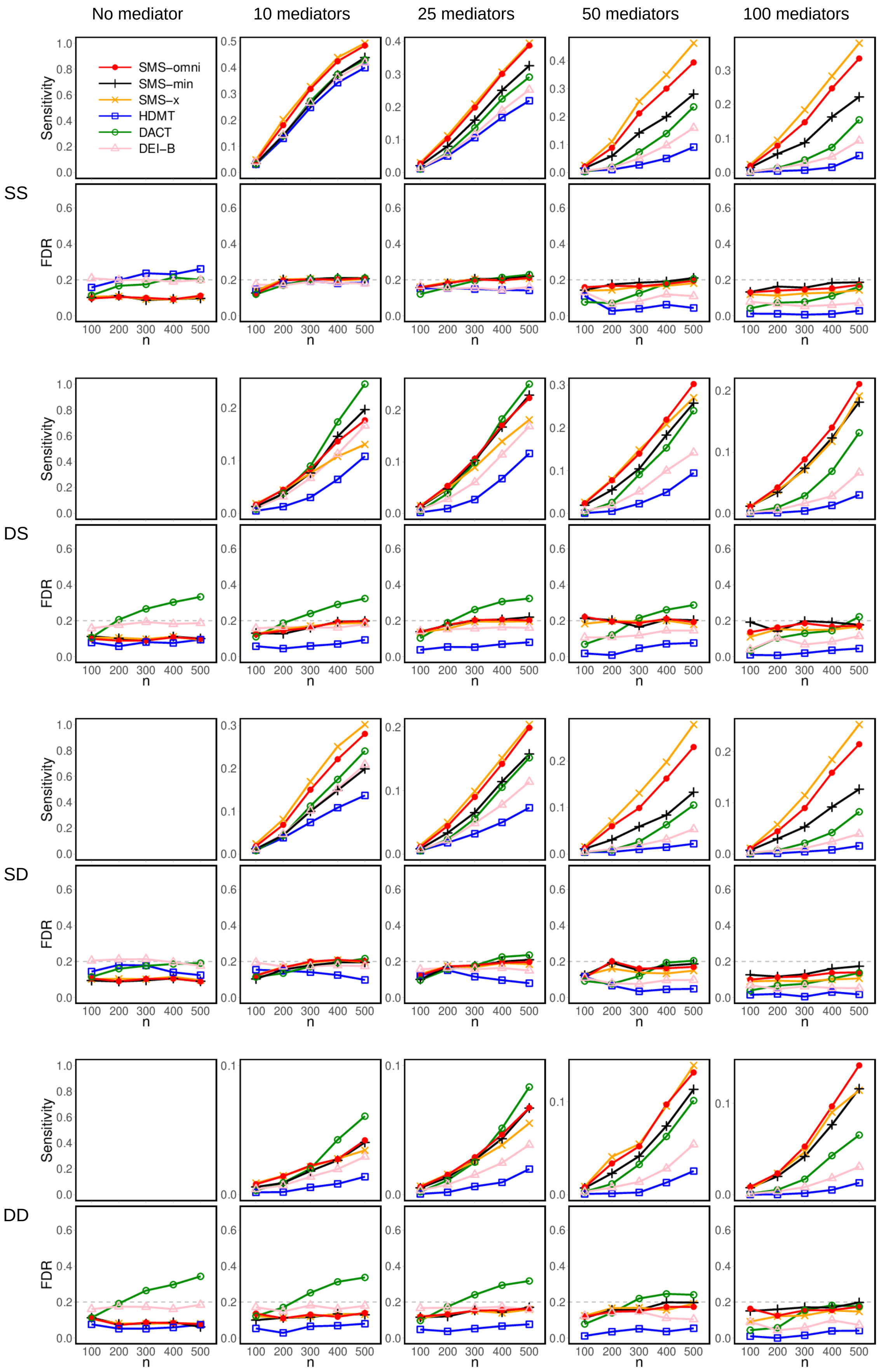




Figure S6: Simulation results on empirical FDR and sensitivity based on $J = 1{,}000$ features and a binary outcome, mirroring the PKU-SPCO metabolomics study. The dashed gray line indicates the nominal FDR level of 0.2.

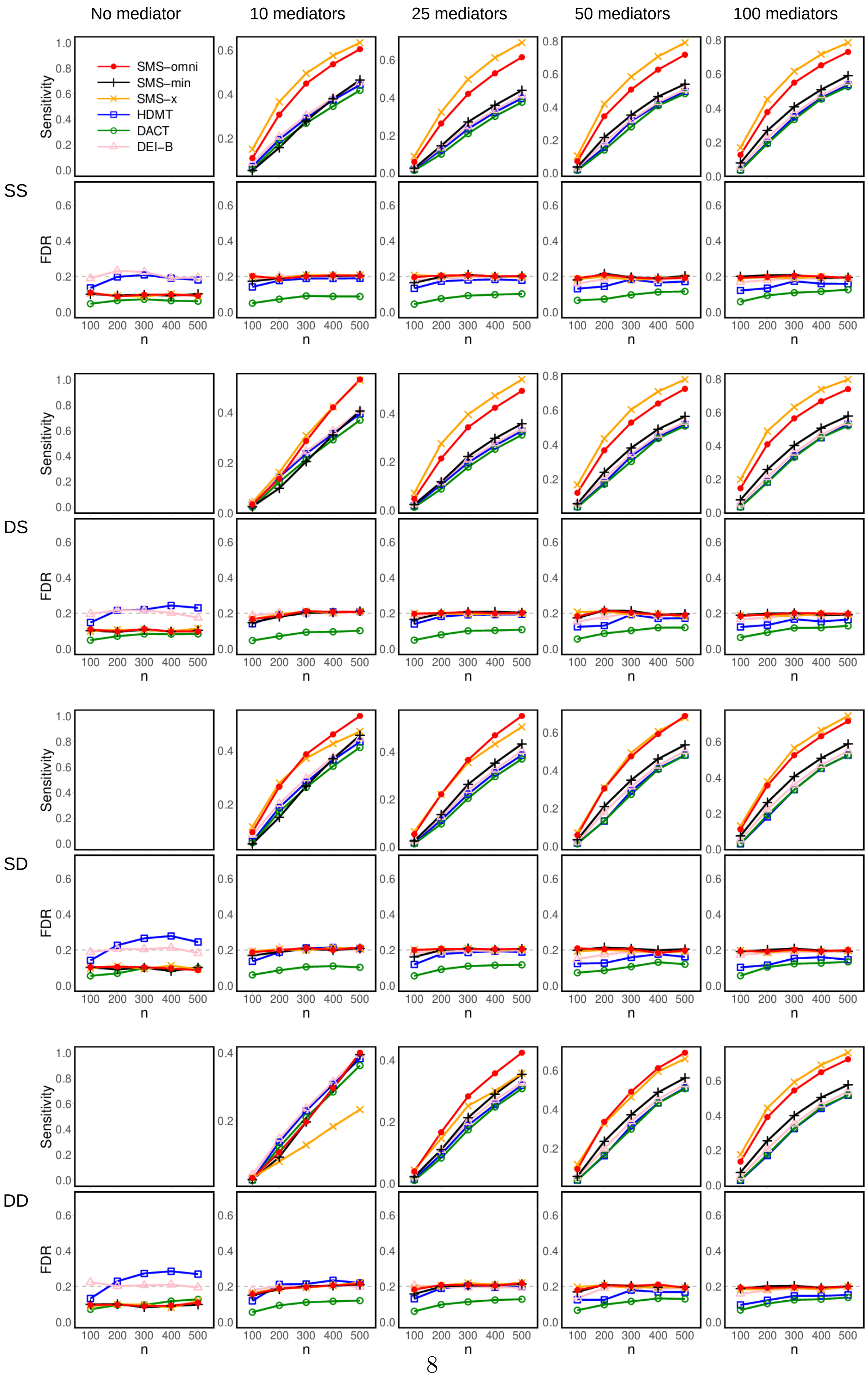




Figure S7: Simulation results on empirical FDR and sensitivity based on $J = 50{,}000$ features and feature-specific continuous outcomes, mirroring the SNP–DNA methylation–gene expression study [7]. The dashed gray line indicates the nominal FDR level of 0.2.

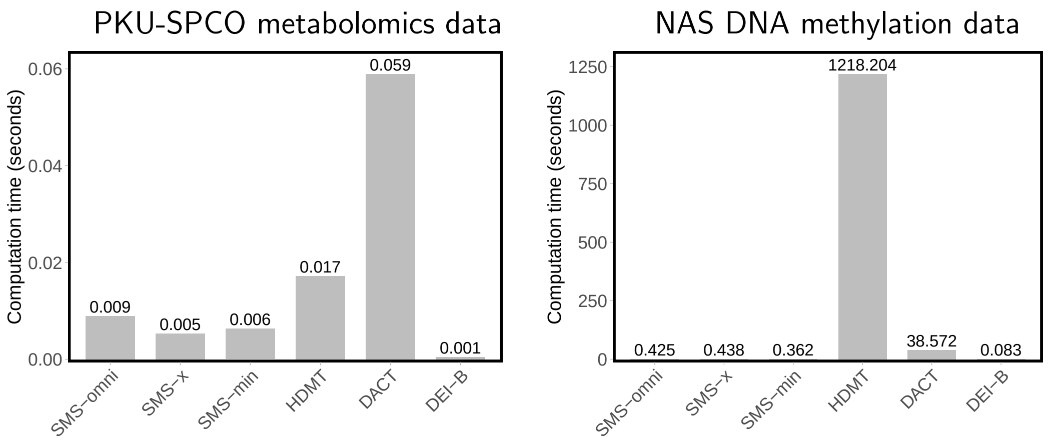


Figure S8: Computation times, measured in seconds, for analyses of the two real datasets. Times were measured after the effect-size estimates, $Z$-scores, and $p$-values for the two associations had been obtained.

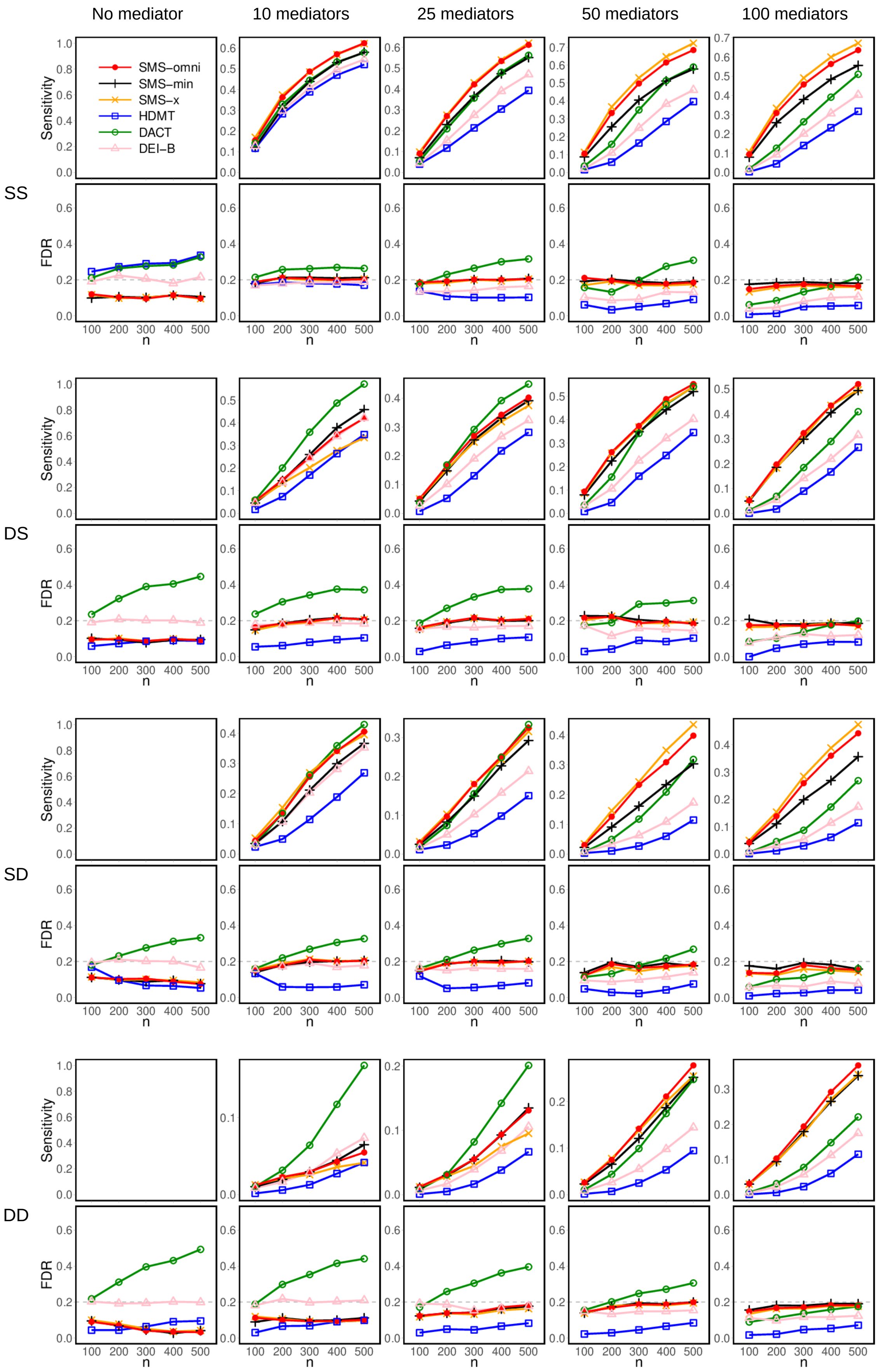




Figure S9: Simulation results on empirical FDR and sensitivity, based on $J = 500$ features. The dashed gray line indicates the nominal FDR level of 0.2.

Table S1: CpG sites identified by SMS-omni in the NAS dataset (FDR = 0.05)

| | | Smoking–CpG methylation association | | CpG methylation–lung function association | | |
|---|---|---|---|---|---|---|
| Rank | CpG site | $Z$-score | $p$-value | $Z$-score | $p$-value | Also identified by |
| 1 | cg05575921 | −8.32 | $5.9 \times 10^{-16}$ | −8.12 | $2.6 \times 10^{-15}$ | HDMT, DEI-B, DACT |
| 2 | cg03636183 | −6.09 | $2.0 \times 10^{-9}$ | −6.43 | $2.5 \times 10^{-10}$ | HDMT, DEI-B, DACT |
| 3 | cg06126421 | −6.66 | $6.4 \times 10^{-11}$ | −5.74 | $1.5 \times 10^{-8}$ | HDMT, DEI-B, DACT |
| 4 | cg21566642 | −6.80 | $2.6 \times 10^{-11}$ | −5.61 | $3.1 \times 10^{-8}$ | HDMT, DEI-B, DACT |
| 5 | cg05951221 | −6.46 | $2.2 \times 10^{-10}$ | −5.17 | $3.2 \times 10^{-7}$ | HDMT, DEI-B, DACT |
| 6 | cg14753356 | −5.24 | $2.3 \times 10^{-7}$ | −4.91 | $1.2 \times 10^{-6}$ | HDMT, DEI-B, DACT |
| 7 | cg23771366 | −5.65 | $2.6 \times 10^{-8}$ | −4.62 | $4.7 \times 10^{-6}$ | HDMT, DEI-B, DACT |
| 8 | cg06644428 | −7.18 | $2.1 \times 10^{-12}$ | −3.78 | $1.8 \times 10^{-4}$ | HDMT, DEI-B, DACT |
| 9 | cg11660018 | −6.43 | $2.7 \times 10^{-10}$ | −3.90 | $1.1 \times 10^{-4}$ | HDMT, DEI-B, DACT |
| 10 | cg01940273 | −5.10 | $4.6 \times 10^{-7}$ | −4.31 | $1.9 \times 10^{-5}$ | HDMT, DEI-B, DACT |
| 11 | cg21322436 | −5.38 | $1.1 \times 10^{-7}$ | −4.17 | $3.5 \times 10^{-5}$ | HDMT, DEI-B, DACT |
| 12 | cg15342087 | −4.25 | $2.5 \times 10^{-5}$ | −5.05 | $5.9 \times 10^{-7}$ | HDMT, DEI-B, DACT |
| 13 | cg24859433 | −4.25 | $2.4 \times 10^{-5}$ | −4.95 | $9.6 \times 10^{-7}$ | HDMT, DEI-B, DACT |
| 14 | cg25189904 | −5.35 | $1.3 \times 10^{-7}$ | −4.01 | $6.9 \times 10^{-5}$ | HDMT, DEI-B, DACT |
| 15 | cg23916896 | −5.13 | $3.9 \times 10^{-7}$ | −4.00 | $7.2 \times 10^{-5}$ | HDMT, DEI-B, DACT |
| 16 | cg25949550 | −4.53 | $7.1 \times 10^{-6}$ | −3.91 | $1.0 \times 10^{-4}$ | HDMT, DEI-B, DACT |
| 17 | cg11902777 | −4.19 | $3.2 \times 10^{-5}$ | −3.95 | $8.6 \times 10^{-5}$ | HDMT, DEI-B, DACT |
| 18 | cg14580211 | −3.26 | 0.0012 | −5.29 | $1.7 \times 10^{-7}$ | HDMT, DEI-B |
| 19 | cg14624207 | −4.04 | $6.1 \times 10^{-5}$ | −3.65 | $2.8 \times 10^{-4}$ | HDMT, DEI-B, DACT |
| 20 | cg03991871 | −4.04 | $6.0 \times 10^{-5}$ | −3.60 | $3.5 \times 10^{-4}$ | HDMT, DEI-B, DACT |
| 21 | cg21161138 | −3.69 | $2.4 \times 10^{-4}$ | −3.82 | $1.4 \times 10^{-4}$ | HDMT, DEI-B |
| 22 | cg11554391 | −3.62 | $3.2 \times 10^{-4}$ | −3.81 | $1.5 \times 10^{-4}$ | HDMT, DEI-B |
| 23 | cg19859270 | −5.03 | $6.5 \times 10^{-7}$ | −3.07 | 0.0023 | |
| 24 | cg09935388 | −5.44 | $7.9 \times 10^{-8}$ | −2.69 | 0.0073 | |
| 25 | cg01692968 | −4.80 | $2.0 \times 10^{-6}$ | −2.81 | 0.0051 | |
| 26 | cg01899089 | −3.76 | $1.8 \times 10^{-4}$ | −3.15 | 0.0017 | |
| 27 | cg10470891 | −3.28 | 0.0011 | −3.48 | $5.4 \times 10^{-4}$ | HDMT, DEI-B |
| 28 | cg13442016 | −3.49 | $5.2 \times 10^{-4}$ | −3.23 | 0.0013 | HDMT, DEI-B |
| 29 | cg05284742 | −4.12 | $4.2 \times 10^{-5}$ | −2.96 | 0.0032 | |
| 30 | cg17247206 | −3.08 | 0.0022 | −3.74 | $2.0 \times 10^{-4}$ | |
| 31 | cg08101174 | −3.55 | $4.2 \times 10^{-4}$ | −3.16 | 0.0017 | |
| 32 | cg25648203 | −3.40 | $7.2 \times 10^{-4}$ | −3.19 | 0.0015 | HDMT |
| 33 | cg13193840 | −3.92 | $9.9 \times 10^{-5}$ | −2.87 | 0.0042 | |
| 34 | cg09880681 | −3.47 | $5.5 \times 10^{-4}$ | −3.01 | 0.0028 | |
| 35 | cg13039251 | 4.96 | $9.2 \times 10^{-7}$ | 2.45 | 0.0147 | |
| 36 | cg06023570 | −3.33 | $9.2 \times 10^{-4}$ | −2.98 | 0.0030 | |
| 37 | cg11279857 | −3.12 | 0.0019 | −3.08 | 0.0022 | |

| | | Smoking–CpG methylation association | | CpG methylation–lung function association | | |
|---|---|---|---|---|---|---|
| Rank | CpG site | $Z$-score | $p$-value | $Z$-score | $p$-value | Also identified by |
| 38 | cg23079012 | −3.12 | 0.0019 | −3.07 | 0.0022 | |
| 39 | cg00008629 | −2.92 | 0.0036 | −3.39 | $7.5 \times 10^{-4}$ | |
| 40 | cg10874644 | 3.14 | 0.0018 | 3.04 | 0.0025 | |
| 41 | cg07845392 | −3.38 | $7.7 \times 10^{-4}$ | −2.90 | 0.0039 | |
| 42 | cg00073090 | −3.37 | $8.1 \times 10^{-4}$ | −2.91 | 0.0038 | |
| 43 | cg17024919 | −3.74 | $2.0 \times 10^{-4}$ | −2.75 | 0.0061 | |
| 44 | cg15723536 | −3.57 | $3.8 \times 10^{-4}$ | −2.79 | 0.0054 | |
| 45 | cg18521743 | −3.54 | $4.3 \times 10^{-4}$ | −2.80 | 0.0053 | |
| 46 | cg25537245 | −2.65 | 0.0083 | −3.96 | $8.4 \times 10^{-5}$ | |
| 47 | cg26963277 | −2.93 | 0.0036 | −3.15 | 0.0017 | |
| 48 | cg01827726 | −3.33 | $9.1 \times 10^{-4}$ | −2.82 | 0.0050 | |
| 49 | cg27447254 | 3.49 | $5.2 \times 10^{-4}$ | 2.75 | 0.0061 | |
| 50 | cg08428878 | 2.94 | 0.0034 | 3.04 | 0.0025 | |
| 51 | cg27303733 | 2.65 | 0.0082 | 3.71 | $2.2 \times 10^{-4}$ | |
| 52 | cg02583484 | −2.94 | 0.0034 | −3.01 | 0.0027 | |
| 53 | cg26077378 | −4.06 | $5.6 \times 10^{-5}$ | −2.53 | 0.0116 | |
| 54 | cg26764244 | −3.79 | $1.6 \times 10^{-4}$ | −2.61 | 0.0092 | |
| 55 | cg12803068 | 3.28 | 0.0011 | 2.80 | 0.0052 | |
| 56 | cg06595693 | −2.60 | 0.0096 | −3.80 | $1.6 \times 10^{-4}$ | |
| 57 | cg00687135 | −3.36 | $8.4 \times 10^{-4}$ | −2.75 | 0.0061 | |
| 58 | cg22937882 | 2.71 | 0.0069 | 3.38 | $7.8 \times 10^{-4}$ | |
| 59 | cg12956507 | −2.95 | 0.0033 | −2.90 | 0.0039 | |
| 60 | cg15857661 | −2.91 | 0.0038 | −2.92 | 0.0036 | |
| 61 | cg05251190 | −2.94 | 0.0034 | −2.90 | 0.0039 | |
| 62 | cg23576855 | −3.12 | 0.0019 | −2.80 | 0.0052 | |
| 63 | cg11231349 | −2.83 | 0.0049 | −3.04 | 0.0024 | |
| 64 | cg05296978 | −2.41 | 0.0163 | −4.18 | $3.3 \times 10^{-5}$ | |
| 65 | cg01901332 | −2.62 | 0.0091 | −3.53 | $4.4 \times 10^{-4}$ | |
| 66 | cg08900404 | −3.59 | $3.6 \times 10^{-4}$ | −2.59 | 0.0099 | |
| 67 | cg13594138 | 3.00 | 0.0028 | 2.82 | 0.0050 | |
| 68 | cg19572487 | −5.08 | $5.2 \times 10^{-7}$ | −2.15 | 0.0318 | |
| 69 | cg25458175 | −3.76 | $1.9 \times 10^{-4}$ | −2.50 | 0.0128 | |
| 70 | cg18900812 | −2.99 | 0.0029 | −2.79 | 0.0054 | |

Note: The CpG sites are ranked in descending order of the SMS-omni statistic.
